# A comprehensive landscape of the zinc-regulated human proteome

**DOI:** 10.1101/2024.01.04.574225

**Authors:** Nils Burger, Melanie J. Mittenbühler, Haopeng Xiao, Sanghee Shin, Luiz H.M. Bozi, Shelley Wei, Hans-Georg Sprenger, Yizhi Sun, Yingde Zhu, Narek Darabedian, Jonathan J. Petrocelli, Pedro Latorre- Muro, Jianwei Che, Edward T. Chouchani

**Affiliations:** Department of Cancer Biology, Dana–Farber Cancer Institute, Boston, MA, USA; Department of Cell Biology, Harvard Medical School, Boston, MA, USA; Department of Biological Chemistry and Molecular Pharmacology, Harvard Medical School, Boston, MA, USA

## Abstract

Zinc is an essential micronutrient that regulates a wide range of physiological processes, principally through Zn^2+^ binding to protein cysteine residues. Despite being critical for modulation of protein function, for the vast majority of the human proteome the cysteine sites subject to regulation by Zn^2+^ binding remain undefined. Here we develop ZnCPT, a comprehensive and quantitative mapping of the zinc-regulated cysteine proteome. We define 4807 zinc-regulated protein cysteines, uncovering protein families across major domains of biology that are subject to either constitutive or inducible modification by zinc. ZnCPT enables systematic discovery of zinc-regulated structural, enzymatic, and allosteric functional domains. On this basis, we identify 52 cancer genetic dependencies subject to zinc regulation, and nominate malignancies sensitive to zinc-induced cytotoxicity. In doing so, we discover a mechanism of zinc regulation over Glutathione Reductase (GSR) that drives cell death in GSR-dependent lung cancers. We provide ZnCPT as a resource for understanding mechanisms of zinc regulation over protein function.

## Introduction

Metal ions are essential micronutrients that play critical roles in all aspects of cellular biology. The zinc ion (Zn^2+^) is among the most widely employed metal cofactors in the cell and the vast majority of cellular zinc is bound to proteins^1,2^. Zinc binds to proteins as a constitutive structural component, to act as a catalyst, or to otherwise regulate target function^1,3^(**Figure 1A**). These interactions are thought to be extremely prevalent, as it is predicted that upwards of 10% of the human proteome could be regulated by zinc binding^4,5^. Within a cell, local zinc concentrations are tightly regulated by zinc transporters as well as zinc storage and carrier proteins, which can drive inducible interactions of zinc with protein targets. As such, zinc binding to proteins is implicated in a wide range of biological processes^1–3^.

**Figure 1:**
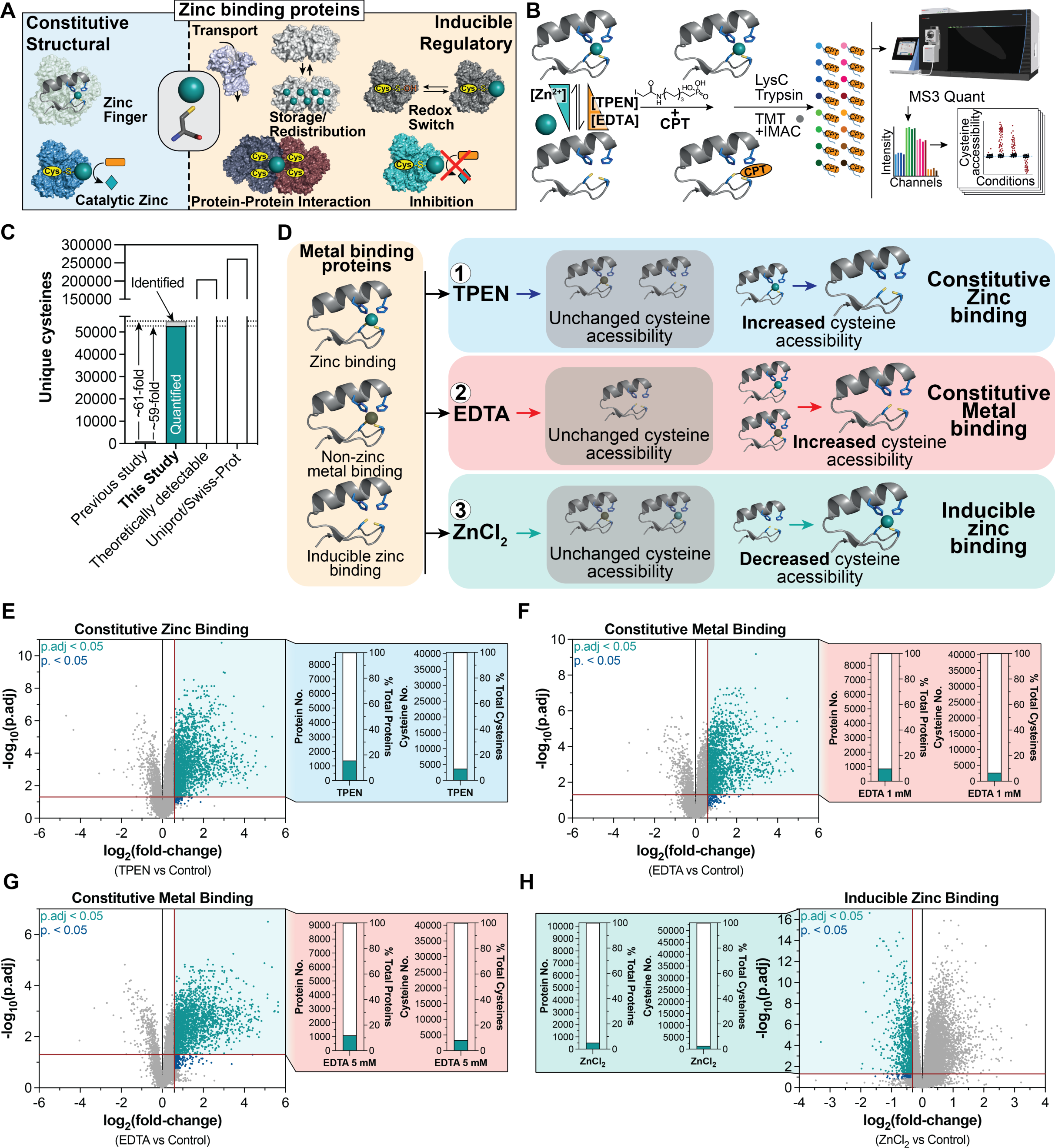
The ZnCPT dataset defines a quantitative map of the zinc binding proteome. A Diverse functions are mediated by protein zinc binding B Chemoproteomic workflow to determine zinc coordination by protein cysteines C Comparison of cysteine coverage of ZnCPT and a previous study^8^ to the theoretically quantifiable cysteine proteome^68^ D Illustration of the three different treatment strategies to determine the constitutive zinc binding, metal binding and inducible zinc binding proteome E Cysteine accessibility changes upon TPEN (1 mM) treatment compared to control F Cysteine accessibility changes upon EDTA (1 mM) treatment compared to control G Cysteine accessibility changes upon EDTA (5 mM) treatment compared to control H Cysteine accessibility changes upon ZnCl_2_ (10 µM) treatment compared to control

Despite the widespread importance of zinc regulation over biological processes, there is a dearth of information regarding the specific protein modifications that explain the mechanistic basis for this activity. To date, identification of zinc-binding sites on proteins has relied on biophysical analyses of individual targets and prediction tools based on conserved sequence features of known zinc-binding proteins^4–6^. Comprehensive proteome-wide analysis of zinc binding to proteins is lacking because of technical challenges due to the non-covalent nature of coordination bonds between protein residues and zinc. Specifically, zinc binding to proteins most frequently involves chelation with at least one cysteine thiol^6,7^, and methods for assessing these modifications on protein cysteine residues to date covered a small proportion of the cysteine proteome^8,9^. For this reason, there has been no systematic mapping of the zinc binding proteome.

Herein we develop a cysteine derivatization and enrichment method coupled with multiplexed proteomics to provide a quantitative and thorough landscape of the zinc-regulated cysteine proteome. This ZnCPT dataset quantifies zinc modification status across over 52,000 cysteines in the human proteome. The zinc modification state of most of these sites had not previously been determined, so this landscape represents by far the deepest examination of the zinc-regulated cysteine proteome.

This compendium allows us to establish and validate distinct zinc-regulated proteins that underlie major aspects of cell biology. From this dataset, we define a structural basis differentiating protein cysteine thiol features that facilitate constitutive binding or inducible binding. We identify distinct clusters of the cysteine proteome constitutively bound by zinc, compared to those, subject to dynamic inducible modification by zinc. In doing so, we identify zinc-regulated structural, enzymatic, and allosteric functional domains on a range of cancer dependencies to nominate malignancies sensitive to zinc-induced cytotoxicity. We discover a mechanism of zinc-driven control over Glutathione Reductase (GSR) that drives cell death in GSR-dependent lung cancer cells. Together, these findings provide a comprehensive analysis of the zinc-regulated human proteome.

## Results

### Cysteine-Reactive Phosphate Tags (CPTs) Provide Deep Coverage and Quantification of the zinc-bound Cysteine Proteome

Zinc binding to proteins most typically involves coordination with at least one cysteine thiolate sidechain^6,7^. Generally, quantification of cysteine thiolate modifications can be determined by cysteine derivatization and quantification approaches^10^. However, the non-covalent nature of cysteine-zinc interactions requires determination of zinc binding to proteins under native conditions to preserve zinc coordination. We were inspired by recent mass spectrometry methodologies that determine zinc binding to protein cysteine residues under native conditions^8,9^. In particular, the elegant strategy developed by Pace & Weerapana^8^ determines modification of protein cysteines by zinc under native conditions. To date, these methods achieve low proteome coverage (∼900 sites, ∼ 0.0034% of the cysteine proteome), due to the low abundance of cysteine residues relative to other amino acids, and a dearth of effective enrichment strategies for cysteine containing peptides. As such, quantification of protein cysteine modification by zinc across the majority of the proteome has been a technical hurdle.

We recently developed an approach to comprehensively identify and quantify the extent of reversible modification of tens of thousands of cysteines across the proteome in a single experiment^10^. This method relies on a cysteine labeling and enrichment reagent for quantitative proteomics, called cysteine-reactive phosphate tags (CPT). CPTs facilitate >99% enrichment of cysteine-containing peptides using metal affinity chromatography (IMAC) enrichment, allowing for unprecedentedly deep quantitative mapping of the cysteine proteome. We posited that CPTs could be deployed to assess quantitative engagement of zinc simultaneously with tens of thousands of cysteines. We devised a strategy combining CPT with tandem mass tag (TMT)-multiplexed chemoproteomics^10,11^, to quantify zinc engagement with over 52,000 unique cysteines across the human proteome **(Figure 1B, C & Supplementary Table 1)**.

We used HCT116 cells as this system captures a large proportion of the human proteome^12,13^, allowing for assessment of zinc engagement with over 10,000 proteins **(Supplementary Table 1).** We treated native HCT116 cell lysates with well-established manipulations to titrate zinc binding to proteins (**Figure 1D; Figure S1A**)^8,14–16^. Following these interventions, we applied a labeling strategy for proteome-wide quantification of zinc binding to protein cysteines, combined with TMT multiplexing^17,18^ that allows for simultaneous analysis of up to 18 biological replicates in a single experiment (**Figure 1B)**.

To define constitutively zinc-bound protein cysteines, we mapped protein cysteine residues that become accessible to CPT modification following treatment with the zinc chelator N,N,N′,N′-tetrakis(2-pyridinylmethyl)-1,2-ethanediamine (TPEN; **Figure 1D)**. In parallel, we defined constitutively metal-bound cysteines by mapping cysteine residues that become accessible to CPT modification following treatment with the broad metal chelator Ethylenediaminetetraacetic acid (EDTA) **(Figure 1D)**. Finally, we determined protein cysteines amenable to inducible zinc modification by defining those that are blocked from CPT labelling following treatment with ZnCl_2_ (**Figure 1D)**. We applied a physiologic concentration of zinc at 10 µM, which is well below total cellular zinc (200-300 µM^2^) and falls near plasma zinc concentration range (11-24 µM^19^). Since local zinc concentrations are dynamic and local spikes in zinc concentration are widely reported, we estimated the concentration used here to be within a conservative physiologic range for inducible zinc binding.

### Population characteristics of the zinc-binding proteome

The vast majority of cysteine sites mapped in our analyses have not been previously experimentally assessed for zinc binding. The major factor contributing to the high proportion of previously unmapped sites is that the CPT method provides over an order of magnitude improvement in cysteine-peptide enrichment compared to previous technologies^10^ **(Figure 1C)**. Of the entire detected cysteine proteome (54,900), 52,665 unique cysteines were quantified.

Global quantification of zinc binding to protein cysteines was remarkably consistent across biological replicates, with the large majority of cysteines exhibiting reproducible extents of modification **(Figure S1B)**. Constitutive zinc- and metal-binding populations clustered closely and distinctly from control and inducible zinc-binding populations. Replicate samples showed extremely high reproducibility of zinc binding quantification across cysteine sites and the same was observed across biological replicates for zinc treated samples **(Figure S2A-E)**.

First, we curated the ZnCPT dataset to define population characteristics of the cysteine proteome that participates in (i) constitutive zinc-binding, (ii) non-zinc metal binding, and (iii) inducible zinc-binding. Of the entire quantified cysteine proteome (52,665), we identified 3,698 constitutively zinc bound cysteines, and 4,328 constitutively metal bound cysteines **(Figure 1E-G)**. Over 10 % of quantified proteins contained at least one constitutive zinc-binding cysteine, which aligns with bioinformatic estimates of the proportion of zinc-binding proteins in the human proteome^4,5^. In addition, we identified 1,358 cysteines that could be inducibly modified by zinc **(Figure 1H)**. Notably, cysteine sites responding to TPEN showed a very similar response upon EDTA treatment, which confirms that zinc is the predominant protein-bound metal coordinated by cysteine residues (**Figure 2A**). In contrast, the number of cysteines presenting with increased accessibility upon EDTA but not TPEN treatment, indicating non-zinc metal binding sites, was comparably low.

**Figure 2:**
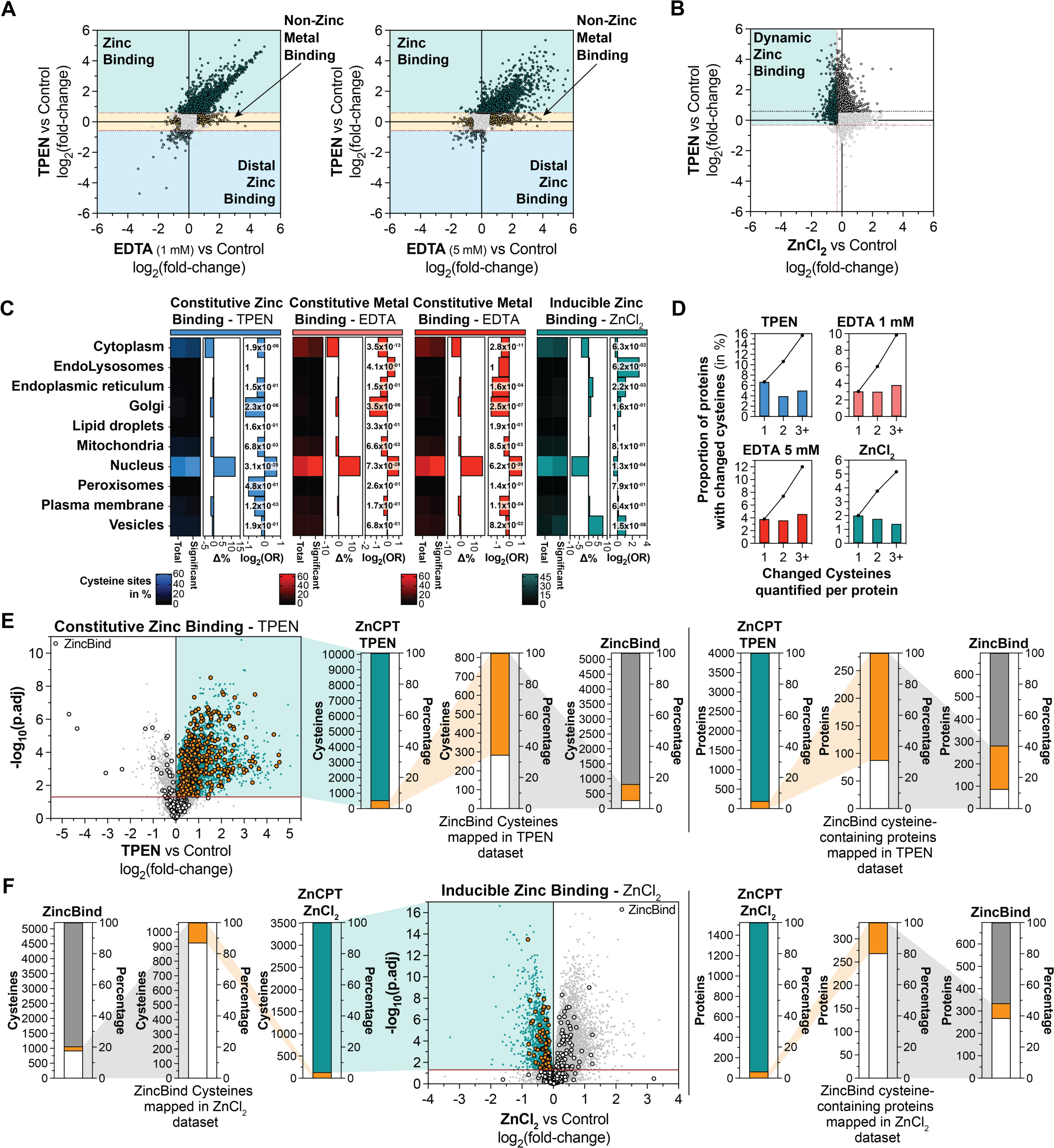
Characterization of zinc and metal binding cysteines. A Comparison of cysteine accessibility changes between EDTA (1 & 5 mM) and TPEN treatment (relative to control) to define metal binding subpopulations B Comparison of cysteine accessibility changes between ZnCl_2_ and TPEN treatment (relative to control) to define dynamic zinc binding cysteine populations C Subcellular distribution and enrichment of all and significantly changed cysteine containing proteins, upon treatment with TPEN, EDTA (1 & 5 mM) or ZnCl_2_ D Distribution of quantified cysteine residues per protein that are significantly changed upon treatment with TPEN, EDTA (1 & 5 mM) or ZnCl_2_ E Comparison of cysteine coverage and cysteine accessibility changes upon TPEN treatment with zinc binding cysteines identified in the ZincBind^6^ dataset. Comparison of coverage of proteins containing cysteines that exhibit accessibility changes upon TPEN treatment to proteins containing zinc binding cysteines identified in the ZincBind dataset. F Comparison of cysteine coverage and cysteine accessibility changes upon ZnCl_2_ treatment with zinc binding cysteines identified in the ZincBind^6^ dataset. Comparison of coverage of proteins containing cysteines that exhibit accessibility changes upon ZnCl_2_ treatment to proteins containing zinc binding cysteines identified in the ZincBind dataset.

As expected, constitutive metal binding cysteine sites were the largest population, which were predominantly populated by constitutive zinc-bound cysteines. Interestingly, a large majority of cysteines amenable to inducible zinc modification represented a completely distinct population of the cysteine proteome (**Figure 2B, S2F, G)**. These data suggest that distinct proximal amino acid environments govern capacity for inducible zinc binding, compared to constitutive zinc binding. The structural basis for this is investigated in a later section. Of note, we also defined a distinct population of cysteines that exhibited increased solvent accessibility as a consequence of zinc addition, indicative of a distal effect of zinc binding, likely induced by structural rearrangements resulting from the zinc binding event. As ZnCPT could not identify the actual zinc binding site in this case, we did not attempt to further investigate cysteines falling into this category, however they are annotated in **Supplementary Table 1**.

We next leveraged the depth of the mapped zinc binding cysteine proteome to generate population-level analyses with subcellular resolution **(Figure 2C)**^20,21^. The constitutive zinc-binding proteome was enriched with nuclear proteins, in line with a significant over-representation of DNA binding proteins (transcription factors, regulators of gene expression) with characteristic structural zinc finger domains. Conversely, there was a de-enrichment of cytoplasmic, Golgi, mitochondrial, and plasma membrane proteins. Interestingly, proteins inducibly regulated by zinc exhibited a distinct subcellular distribution. Inducible zinc targets were de-enriched in the nucleus, and were instead found predominantly vesicular proteins and proteins annotated to be localized within the Endo-/ Lysosomal system, as well as the endoplasmic reticulum. Together, these data indicate that a substantially distinct proteome is targeted by dynamic zinc binding when compared to constitutive zinc-binding proteins. 15.7% of all detected proteins contained at least one cysteine residue that was constitutively modified by zinc while fewer than 5% carried three or more highly modified sites (**Figure 2D, S2H**). For inducible zinc binding, we observed that more than 5% of proteins contained at least one cysteine that was dynamically modified by zinc (**Figure S2H**).

### ZnCPT recapitulates the established zinc-binding proteome

We next examined if ZnCPT recapitulated the cumulative historically determined zinc binding proteins found in the literature. We selected the ZincBind dataset^6^, a compendium of all available structures of zinc binding proteins, as the to-date most comprehensive experimentally validated dataset. Overall, ∼69 % of previously determined zinc binding proteins were reproduced by the ZnCPT constitutive binding dataset (TPEN) **(Figure 2E)**. Conversely, a further ∼20 % were recapitulated by the ZnCPT inducible zinc binding dataset (ZnCl_2_) **(Figure 2F)**. Together, the entire ZnCPT dataset recapitulated ∼70 % of historically accumulated evidence of zinc binding proteins. While ZnCPT reproduced most previously observed zinc binding proteins, some were not recapitulated in this dataset. Some likely reasons for lack of complete overlap include the possibility that conditions used to determine zinc binding of recombinant proteins during structural determination may not recapitulate in a native cellular environment. It is also possible that removal of zinc from proteins in ZnCPT may result in structural rearrangements or aggregation of some proteins which may preclude their analysis by ZnCPT.

Importantly, fewer than 5 % of zinc binding cysteines identified by ZnCPT have been denoted by the ZincBind dataset **(Figure 2E, F)**. This indicates that the ZnCPT compendium substantially expands the experimentally validated cysteine proteome. Furthermore, most inducible zinc binding proteins were not found in ZincBind, underlining the potential of ZnCPT as powerful resource to define the largely undiscovered realm of dynamic zinc binding proteins.

### ZnCPT accurately reports zinc regulation of established zinc-binding proteins

Individual analysis of the protein targets of zinc binding allowed us to investigate modes of zinc regulation over established zinc-binding proteins. We observed many examples of known zinc binding proteins that ZnCPT classified as constitutively bound by zinc. One prominent example is zinc-finger protein 1 (ZPR1), a highly conserved regulator of growth factor signaling via receptor tyrosine kinases, cell proliferation, and translational regulation^22–24^. Based on crystal structures, ZPR1 contains two C4 zinc finger domains (**Figure 3A**). We mapped 6 of these cysteine residues (C80, C83, C259, C262, C288 and C291) as constitutive zinc binding cysteines that could not be further modified by exogenous zinc, indicating full zinc occupancy of all cysteine sites. Another example is the NDUFS6 subunit of mitochondrial complex I which contains a structural zinc finger domain and is critical for assembly of complex I and highly conserved (**Figure 3B**)^25–27^. The zinc is coordinated by three cysteines and one histidine residue, and we quantified complete constitutive zinc binding at these sites with no changes upon zinc treatment.

**Figure 3:**
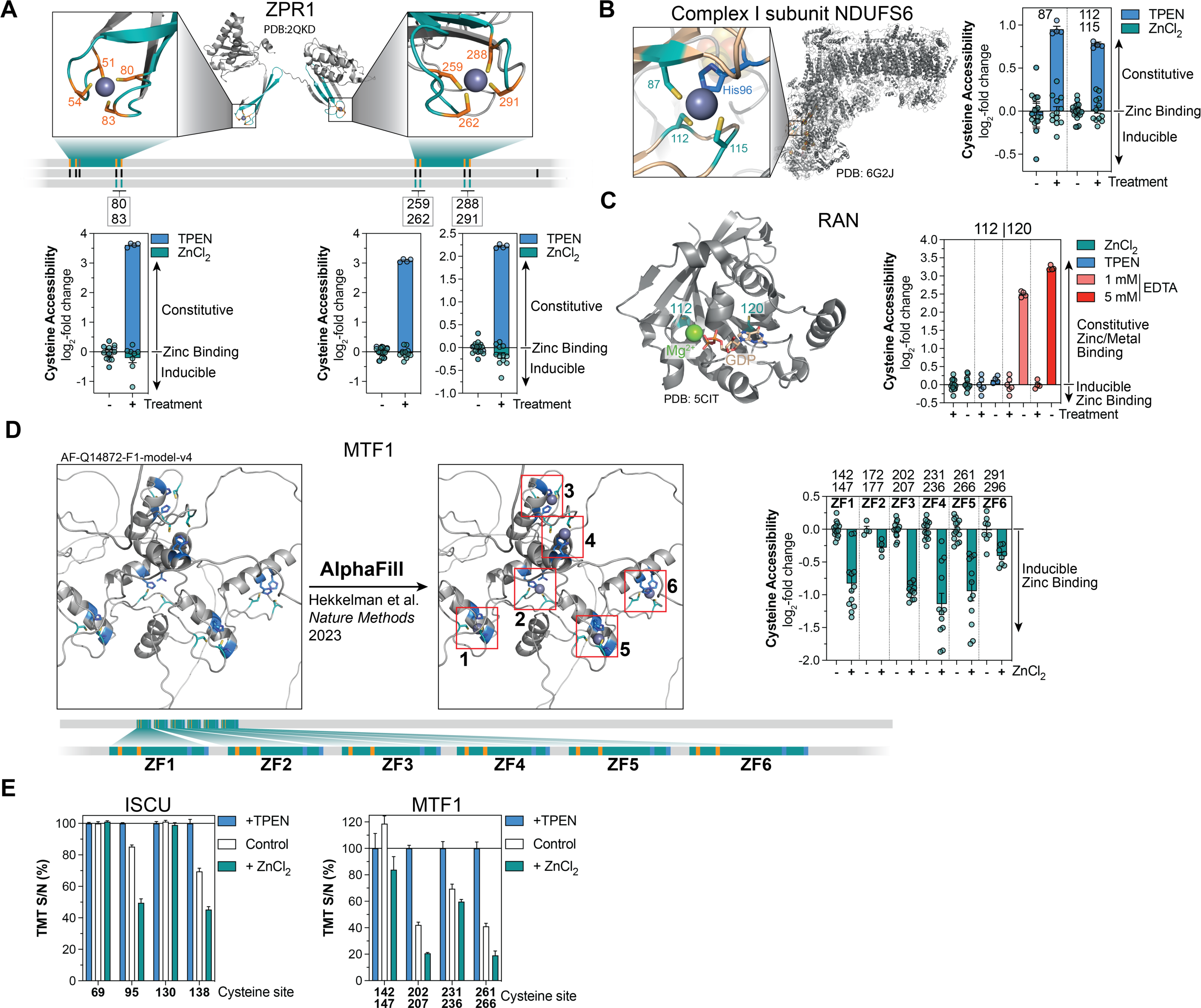
ZnCPT replicates established examples of constitutive and inducible zinc/metal binding. A Accessibility changes of quantified zinc-finger constituting cysteines (Cys80, 83, 259, 262, 288, 291) in ZPR1 upon TPEN/ZnCl_2_ treatment B Accessibility changes of quantified zinc-finger constituting cysteines (Cys87, 112, 115) in the NDUFS6 subunit of mitochondrial complex I upon TPEN/ZnCl_2_ treatment C Accessibility changes of quantified cysteines (Cys112, 120) in RAN, which are occluded by Mg^2+^ and GDP, upon TPEN/EDTA/ZnCl_2_ treatment D Structure of MTF1 predicted by AlphaFold^69^, prior to and post addition of zinc using AlphaFill^33^. Accessibility changes of quantified zinc finger constituting cysteines in MTF1 upon ZnCl_2_ treatment E Quantification of cysteine peptides upon TPEN/ZnCl_2_ treatment compared to control, demonstrates dynamic zinc binding by select cysteines in ISCU and MTF1

ZnCPT also identified numerous examples of known non-zinc metal-binding cysteines. For example, the small GTPase Ran, binds a Mg^2+^ ion that is required for stabilizing GDP/GTP binding^28,29^. The two cysteines of Ran quantified by ZnCPT are occluded by a Mg^2+^ ion which helps to coordinate the phosphate groups of a guanine nucleotide (**Figure 3C**). EDTA, but not TPEN, significantly increased the accessibility of both cysteines, presumably due to its chelation of the Mg^2+^ ion and the resulting destabilization of the nucleotide binding, resulting in the exposure of both cysteine residues (**Figure 3C**). From this we concluded that ZnCPT classification correctly identifies known non-zinc metal binding protein targets which contain cysteine residues, that are occluded as consequence of non-zinc metal coordination.

An example of dynamic zinc binding is the metal binding transcription factor MTF1, which ZnCPT identified as containing six relatively low affinity zinc finger domains to sense free unbound zinc within the cell (**Figure 3D**)^30^. Elevated zinc levels result in the dynamic binding and stabilisation of MTF1 zinc-finger domains^31,32^, facilitating its binding to metal response elements (MREs), driving the transcriptional response to elevated zinc^30^. AlphaFill^33^ derived MTF1 structures identified the dynamically regulated cysteines determined by ZnCPT to be those constituting the six zinc finger domains (**Figure 3D**). These data confirm the validity of the ZnCPT approach to measure occupancy of dynamic zinc-regulated sites. Generally, we found that under control conditions, many zinc-inducible sites exhibited partial zinc occupancy and could be further occupied upon ZnCl_2_ treatment. Importantly, even upon treatment with 10 µM ZnCl_2_ we did not fully occupy dynamically zinc-regulated sites in ISCU (detailed further beneath) and MTF1, confirming that the chosen concentration is in a physiologically relevant range for zinc regulation (**Figure 3E**). Together, ZnCPT provides potential in the identification and characterisation of physiologically relevant dynamic zinc-mediated regulatory events.

### Structural and sequence features of the zinc-binding proteome

ZnCPT comprised the identification of thousands of novel constitutive and inducible zinc binding sites across the proteome. As a first step, we defined protein domains subject to zinc binding by mapping Pfam domain annotations to ZnCPT. This identified classic zinc binding domains as highly enriched, with C2H2 type zinc finger domains dominating (**Figure 4A**) and demonstrated accuracy of our approach in defining zinc binding on a cysteine site level. Next, we interrogated primary sequence motifs (+/- 6 positions of quantified cysteine) to identify primary amino acid signatures as determinants for zinc binding. Highly conserved elements were identified such as vicinal cysteine doublets involved in zinc coordination and certain amino acid nearby that may play important structural roles such as glycine (**Figure 4B, S3A**). Notably, similar amino acid signatures were also identified for inducible zinc binding sites, with additional unique features (**Figure 4C, S3B**). For instance, the prominent lysine residues in inducible binding compared with constitutive site potentially indicates that local electrostatic interactions could be relevant for zinc binding to cysteine thiolates, whereas acidic residues might contribute to zinc coordination and prevent cysteine oxidation.

**Figure 4:**
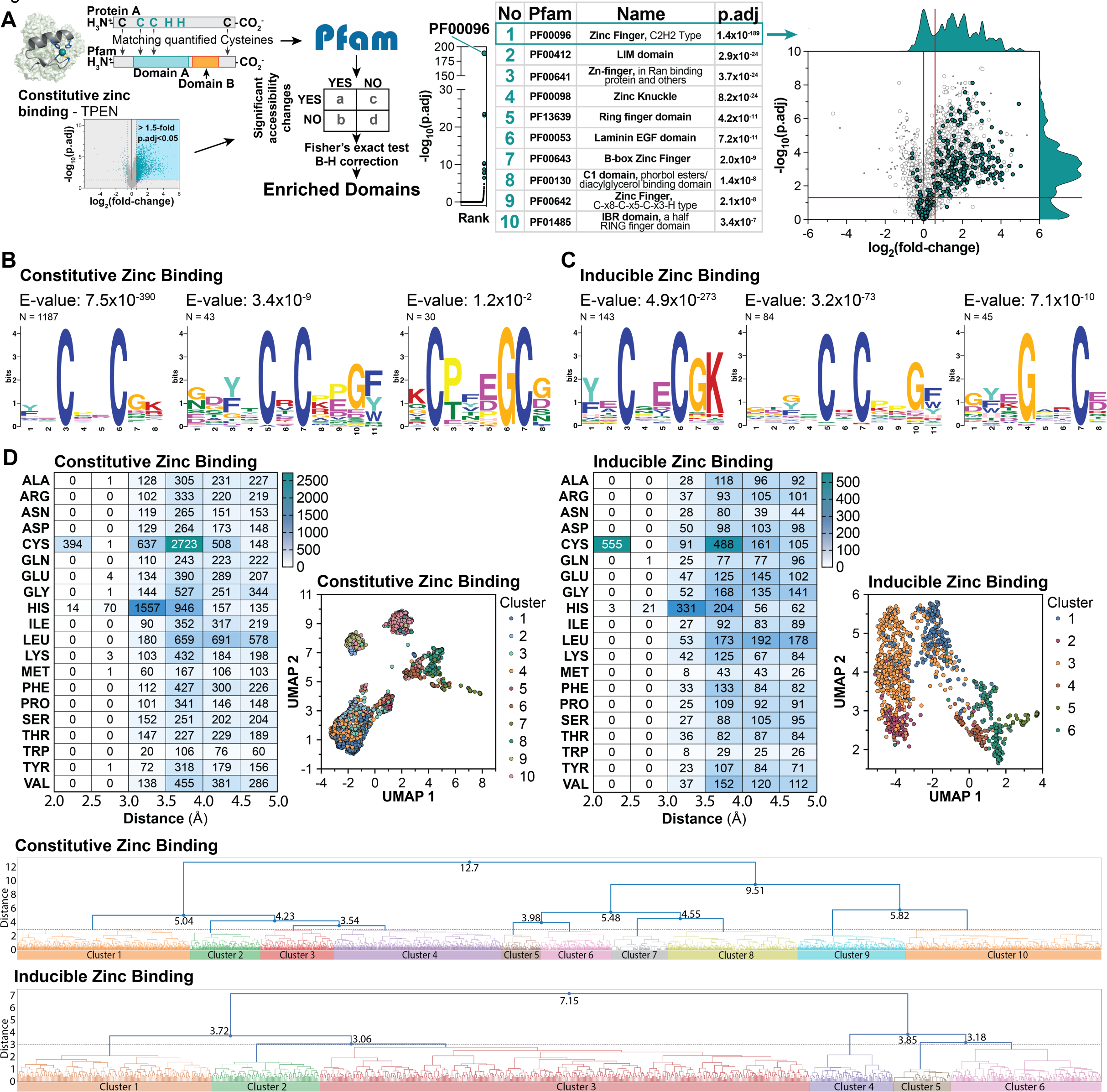
Primary sequence and structural features determine constitutive and inducible zinc binding. A Pfam domain enrichment for constitutive zinc binding cysteines B Primary sequence motifs of constitutive zinc binding cysteine sites C Primary sequence motifs of inducible zinc binding cysteine sites D Distance matrix determining amino acid abundance and distance from inducible or constitutive zinc binding cysteines in protein structures obtained from AlphaFold^69^. Structures of zinc binding sites were clustered based on proximity of surrounding amino acid residues and are additionally illustrated as UMAP plots.

We next examined structural characteristics defining zinc coordination in three dimensions, across the ZnCPT dataset. We systematically analyzed ZnCPT cysteines using human AlphaFold^69^ (AF) structures (https://alphafold.ebi.ac.uk/). This analysis defined the protein microenvironments of cysteines in zinc binding sites (**Figure 4D**, see methods for details). Consistent with primary sequence motif analysis, some general features were readily found s distinct or shared across constitutive and inducible. The presence of nearby cysteine residue(s) and histidine is apparent in both types of zinc binding. This is due to typical C2H2, C3H, and C4 zinc coordination structures in proteins. On the other hand, disulfide bonds suggested by close distance of cysteines (<3 Å) was much more frequent in inducible sites than constitutive sites.

As 3D structures capture coordinating residues distal in sequence space that are impossible to be identified from short motif analysis, unbiased clustering of the site 3D microenvironment was performed. The hierarchical structure clearly identified structurally distinct classes for both constitutive and inducible sites, with 10 clusters for constitutive sites and 6 clusters for inducible sites (**Figure 4D, S3C, D**). For example, amongst constitutive zinc binding sites, cluster 1 and 4 mostly represented single cysteine sites often embedded by hydrophobic residues, and cluster 2 was dominated by disulfide structures. It should be noted that disulfide structures predicted by AF could be in fact more dynamic than a static stable state. Certain structures with two disulfide bonds next to each other could form C4 coordination for Zn^2+^. Cluster 5 represented the more traditional C2H2 type, while there were many members in cluster 6 representing C3H coordination. Interestingly, cluster 7 and 8 also contained mostly C2H2 but with a basic residue such as lysine or arginine and an acidic residue immediately adjacent to the binding site for cluster 7 and an acidic residue for cluster 8. The relationship can also be clearly seen in the UMAP plot where cluster 6-8 were clustered together and away from others. Cluster 9 and 10 were dominated by C4 and C3H, respectively. For inducible zinc binding sites, C3H & C4 were highly represented by cluster 4, whereas C2H2 were represented by clusters 5 and 6. There were also cases of 6 proximal cysteines in cluster 4, presumably coordinating 2 Zn^2+^ to form a Zn_2_C_6_ configuration. These clusters were distant from other clusters in the UMAP plot (**Figure 4D**), indicating potentially different structural mechanisms. In cluster 1, the zinc binding site often involved a single cysteine accompanied by an acidic residue and histidine, where water molecules or the acidic side chain could become the additional coordinating partner upon Zn^2+^ binding. In contrast, clusters 2 and 3 were dominated by disulfides. Taken together, this comprehensive analysis provided a structural basis for the diversity of zinc binding sites including physicochemical determinants of zinc coordination.

### The zinc binding cysteine proteome is distinct from the redox-regulated cysteine proteome

We found that zinc coordinating cysteines are defined by characteristic structural features that markedly differ from features that are characteristic for redox regulated cysteines^10^. These data suggest that zinc-regulated and redox-regulated cysteines are predominantly distinct populations. To examine this, we compared the zinc binding proteome with the recently determined landscape of cysteines subject to reversible redox regulation^10^. Approximately 12,000 unique cysteines were shared across the datasets and conserved across species (human and mouse). Remarkably we observed that cysteines subject to high degrees of redox modification were largely absent from both the constitutive and inducible zinc binding cysteine proteome, albeit the overlap of the redox dataset was proportionally lower for zinc-regulated cysteines compared to non-regulated **(Figure 5A)**. This suggests that cysteine oxidation and zinc binding target distinct populations of the cysteinome. Notably, zinc-binding cysteine motifs are devoid of a proximal arginine residue^10^, a characteristic feature of redox regulated cysteines, instead acidic as well as lysine residues mark zinc-binding cysteines **(Figure 4B-D)**. Together, this implies that inducible zinc binding might regulate target cysteines less amenable to oxidation.

**Figure 5:**
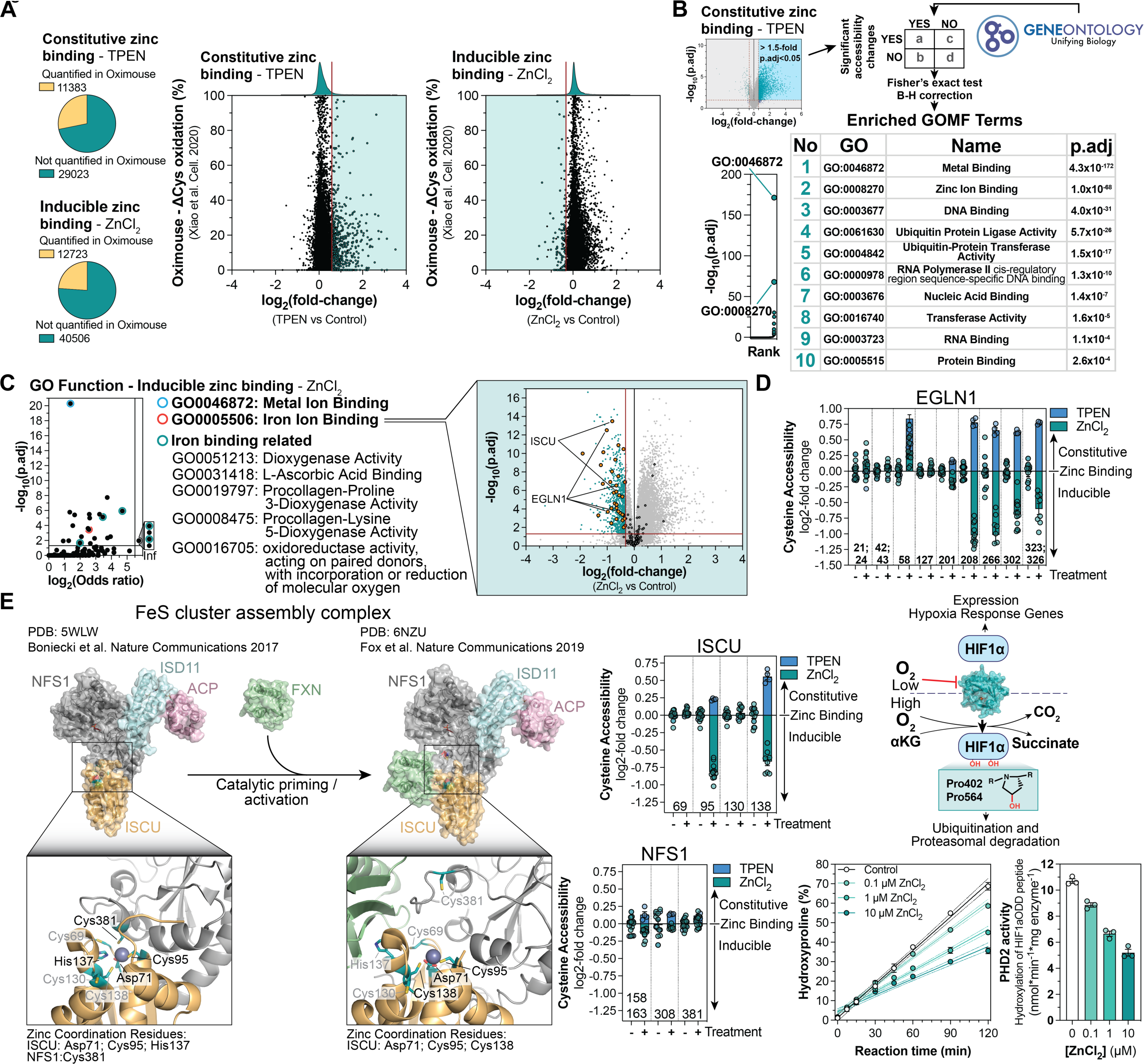
Zinc regulates diverse functional protein classes including iron binding proteins. A Comparison of cysteine oxidation (Oximouse dataset^10^) with accessibility changes upon TPEN/ZnCl_2_ treatment B GO Term (Function) enrichment for constitutive zinc binding cysteines C GO Term (Function) enrichment for inducible zinc binding cysteines identifies iron binding proteins as enriched D The HIF1α prolyl hydroxylase EGLN1 (PHD2) is dynamically regulated by zinc (TPEN/ZnCl_2_). Zinc inhibits its hydroxylation activity E The Iron-Sulfur (FeS) cluster assembly complex can coordinate zinc by ISCU Asp71, Cys95, His137 and NFS1 Cys381. Upon association with Frataxin (FXN), the zinc coordination rearranges and is constituted by ISCU Asp71, Cys95 and Cys138. Cysteine accessibility changes upon TPEN/ZnCl_2_ treatment of quantified cysteines in ISCU and NFS1 are shown.

### Zinc regulates iron-binding proteins

The large majority of zinc-bound cysteine sites in the ZnCPT dataset had not been determined previously. As such, we next sought to systematically classify the biological activities of the zinc-binding proteome. Strikingly, constitutive and inducible zinc binding targets coalesced to largely distinct biological functions **(Figure 5B**, C, Supplementary Table 2). Protein functions related to nucleic acid and ubiquitin binding were prominently enriched amongst constitutive zinc-binding proteins **(Figure 5B)**. To our surprise, proteins determined inducible zinc binding appeared enriched for iron binding protein classes, in addition to others **(Figure 5C, Supplementary Table 2)**. Notably, metal binding to proteins is governed by the individual ligand affinity of metals, as defined in the Irving-Williams series and cellular metal concentrations^35,36^. Changes in metal homeostasis can therefore result in alternative metalation events. In this context, iron displacement by zinc is an established feature, directed by the superior ligand affinity of zinc over iron^36,37^. In total ZnCPT mapped 14 iron binding proteins as inducible zinc binding targets, amongst which iron-dependent dioxygenases predominated. This protein class is exemplified by EGLN1 (PHD2), a principal component of the HIF1α signaling pathway. The enzyme hydroxylates two proline residues within the oxygen-dependent degradation domains (N-terminal NODD and C-terminal CODD) of HIF1α, thereby marking its substrate for ubiquitin-dependent degradation^38,39^. EGLN1 featured pronounced dynamic zinc binding at cysteines known to coordinate catalytic iron **(Figure 5D)**. Indeed, using human recombinant EGLN1, we determined a concentration-dependent inhibition of prolyl hydroxylation on a synthetic peptide resembling part of the human HIF1α-CODD domain, upon ZnCl_2_ treatment **(Figure 5D)**. These data support an inhibitory mechanism of zinc dependent iron displacement for this target.

ZnCPT also provided systematic insights into modes of dynamic zinc regulation over iron-binding proteins, for example the mitochondrial *de novo* iron-sulfur cluster (ISC) assembly machinery^40–44^. Zinc has been implicated as an important cofactor for the ISC assembly machinery, but its physiological role remained obscure^40,43–50^. Zinc can be coordinated by ISCU residues Cys95, Asp71 and His137 and NFS1 Cys381 (**Figure 5E**)^40,44^. Binding of ISC assembly activator Frataxin (FXN), displaces the NFS1 loop carying Cys381 and reorients zinc coordination to ISCU residues Cys95, Asp71 and Cys138^42,43,51,52^. This possibly primes the complex for a catalytic cycle^42,43,53,54^. Importantly, iron, but not zinc binding allowed FeS formation by an *in vitro* ISC assembly complex^53^. ZnCPT quantified both ISCU Cys95 and Cys138 as dynamic zinc binding residues, while NFS1 Cys381 did not appear to bind zinc, supporting a Frataxin-bound intermediary state, awaiting initation of catalysis (**Figure 5E**)^43,44,55^. These data provide physiological evidence of dynamic zinc-dependent regulation of ISC assembly in human cells, which possibly also extends to the wider family of ISC binding proteins^56^.

### Systematic functional classification of zinc-regulated oncoproteins

We reasoned that mapping zinc binding cysteines onto proteome networks can reveal tandem cysteine zinc modifications that regulate proteins’ shared biological activities. Using enrichment analyses for KEGG pathways we identified numerous protein networks and protein pathways dominated by zinc-regulated proteins, both constitutive and inducible **(Figure 6A, S4A)**. For instance, ZnCPT correctly identified all constitutive zinc binding subunits of RNA polymerase except for one, highlighting precision and ultra-deep coverage as strength of this technology (**Figure S4A, B**). Furthermore, Ubiquitin-mediated proteolysis was highly enriched, with ZnCPT identifying E1 SUMO-activating enzymes and E3 Ubiquitin/SUMO protein ligases as constitutive zinc binding, in agreement with the evolutionary conservation of zinc finger domains in ubiquitin/SUMO binding proteins^57,58^(**Figure S4A, C**). Together, these analyses provide a repertoire of functional zinc targets that can be leveraged to define zinc regulation over cellular physiology and to target zinc-dependent pathways in disease.

**Figure 6:**
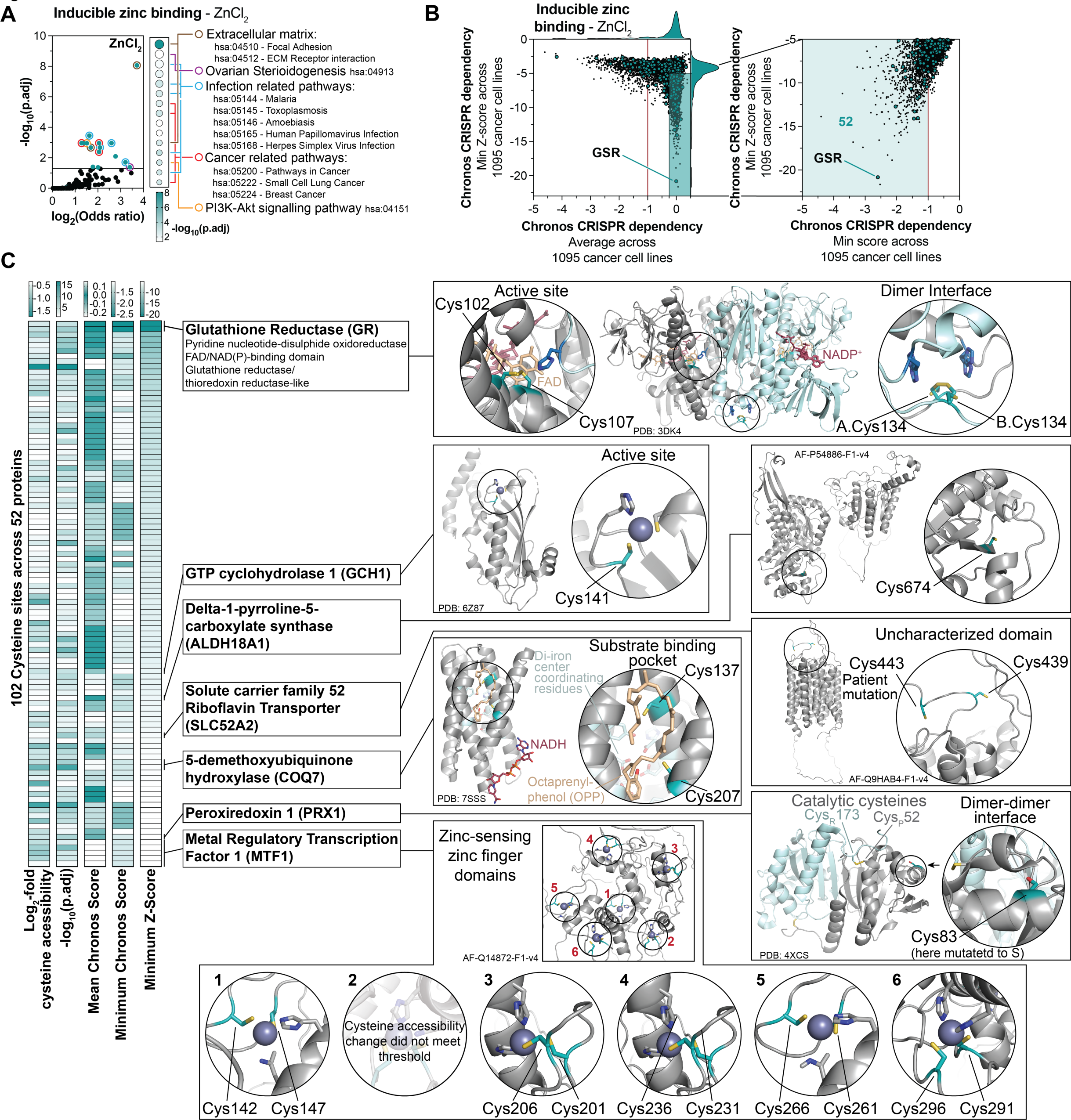
Pathway analysis establishes link between zinc binding and cancer. A KEGG Pathway enrichment for inducible zinc binding cysteines (ZnCl_2_) identifies cancer related pathways as zinc-regulated. B Mapping the Chronos CRISPR dependency score (DepMap Consortium) against the minimum Z-score defines populations of essential genes and selective cell line specific dependencies amongst inducible (ZnCl_2_) zinc binding proteins. Targets with average Chronos CRISPR dependency > −0.25 and minimum Z-score < −5 are selected and the minimum Z-score is mapped against the minimum Chronos CRISPR dependency score to identify strong selective dependencies (minimum Chronos CRISPR dependency < −1). Glutathione Reductase (GSR) has the second highest selective dependency and is identified as target of inducible zinc binding. C ZnCPT identifies numerous selective cancer dependencies defined in DepMap as targets of dynamic zinc regulation, covering diverse functional protein domains

Among the most enriched zinc-regulated proteins we discovered were those involved in regulating tumorigenic processes. In fact, many well established cancer drivers and cancer dependencies were identified as zinc-regulated by ZnCPT (Figure 6A, S4A, D). The discovery of numerous cancer-related proteins being subject to engagement by zinc provided an opportunity to nominate cancer dependencies that could be subject to therapeutic regulation by zinc. To examine this idea systematically, we combined the ZnCPT dataset with DepMap, a comprehensive collection of genetic dependencies across 1095 cancer cell lines. To identify both common essential cancer dependencies, as well as cancer cell-specific dependencies, we ranked average Chronos Cancer Dependency scores and minimum Z-scores **(Figure S5A)**. Correlating both scores denoted two major subpopulations that are zinc-regulated: general essential proteins, and selective cancer dependencies **(Figure 6B, S5B)**. From this analysis, we identified 123 constitutive and 52 inducible zinc-regulated major selective genetic dependencies in DepMap. We posited that targeting zinc regulation might be a powerful strategy to manipulate the function of these cancer dependencies, which could regulate therapeutic response in these cancers.

To further explore this idea, we curated all inducible zinc-regulated cancer dependencies, identifying numerous zinc-regulated functional domains of established cancer dependencies such as Glutathione Reductase (GSR), GTP cyclohydrolase 1 (GCH1), Delta-1-pyrroline-5-carboxylate synthase (ALDH18A1), Riboflavin Transporter (SLC52A2), 5-demethoxyubiquinone hydroxylase (COQ7), Peroxiredoxin 1 (PRX1), and Metal Regulatory Transcription Factor 1 (MTF1) **(Figure 6C)**. From this analysis we selected Glutathione Reductase (GSR) as the most prominent zinc-regulated selective cancer dependency **(Figure 6B, C).** In particular, we found that among lung cancers, SKMES1 lung cancer cells display by far the highest selective GSR dependency **(Figure S5C)**. To gain a deeper understanding of the potential role of GSR in lung cancer, we assessed the expression and protein abundance^59^ of GSR across 685 cancer cells lines and determined a strong correlation **(Figure S5D)**. Importantly, we discerned an elevation in GSR expression in lung compared to other cancer cell lines, which was driven by a population of high GSR-expressing lung cancer types. Notably, these cancers also exhibited elevated expression of GCLC and GCLM, which catalyze the rate-limiting reaction of GSH synthesis, as well as G6PD and PGD, both central NADPH-producing enzymes of the pentose phosphate pathway **(Figure S5D)**. These findings were complemented by a relative elevation of GSH, GSSG and NADP levels^60^, suggesting that glutathione redox metabolism is critical for GSR^High^ lung cancers, possibly to protect against elevated oxidative stress. This motivated us to further elucidate the pronounced relation between GSR and select lung cancers as functional targets of zinc.

### GSR is a target of dynamic zinc binding and zinc-induced cytotoxicity in select non-small cell lung cancers

GSR is a critical enzyme for maintaining the cellular redox homeostasis by keeping the glutathione (GSH) pool reduced. Glutathione metabolism has been widely studied as a therapeutic vulnerability in numerous cancers^61,62^, and our findings suggested a particular importance for GSR metabolism in lung cancer. GSR is a functional homodimer, with its active site constituted by both subunits (**Figure 7A, C**). The enzyme contains two active site cysteine residues that catalyze the reduction of oxidized glutathione (GSSG) into 2x GSH ^63,64^. ZnCPT quantified six cysteines, including the two active site cysteines (Cys102 and Cys107), in GSR, among which the active site and the dimer interface were targets for inducible zinc binding (**Figure 7B, C**). We investigated the functional consequences of zinc binding to GSR and found that zinc can act as potent inhibitor of human recombinant GSR with an IC_50_ of below 1 µM (**Figure 7D**). Furthermore, we observed a slight increase in GSR activity upon titration of TPEN, suggesting a low-level inhibition of enzyme activity by zinc at baseline (**Figure 7D**). To designate the target site of inhibitory zinc binding, we generated recombinant C134S mutant of human GSR and found that there was no perturbation of zinc-mediated enzyme inhibition (**Figure 7E**), indicating the active site of GSR was the key functional target of zinc. These results were corroborated by isothermal calorimetry (ITC) measurements which established a K_d_ of 5.2 +/- 2.6 µM for the wild-type enzyme (**Figure 7F**). In contrast, replacing the active site histidine for a lysine (H511K, catalytic dead mutant) and thereby disrupting the putative zinc binding site in the catalytic domain of the enzyme, reduced the binding affinity for zinc by two orders of magnitude (**Figure 7F**). This data supports zinc-mediated inhibition of GSR by binding to its active site. To examine how Zn^2+^ could be incorporated, we applied the mixed quantum mechanics and molecular mechanics (QM/MM) calculations to optimize a structural model with Zn^2+^ present at the active site. The energy minimized model suggested that tetrahedral coordination provided by Cys102, Cys107, Thr383, and His511 from the partner chain could be stable, consistent with the mutation data (**Figure 7G**).

**Figure 7:**
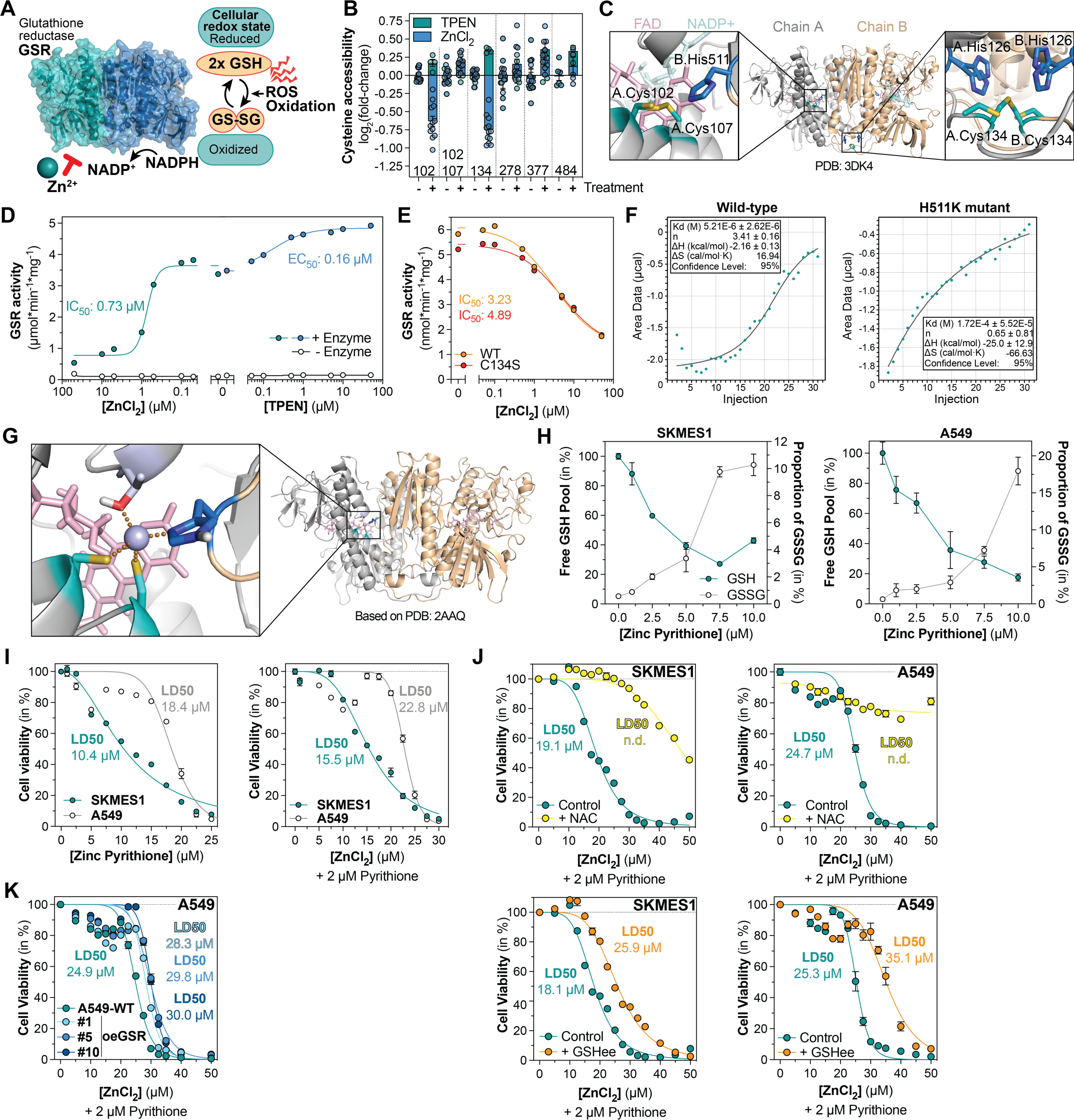
ZnCPT identifies glutathione reductase (GSR) as cancer vulnerability targetable by zinc. A Glutathione Reductase (GSR) is a critical regulator of the cellular redox state and functional dimer utilizing NADPH as redox equivalent to reduce GSSG into GSH. B Accessibility changes of quantified cysteines in GSR upon TPEN/ZnCl_2_ treatment C Possible coordination sites for zinc in GSR are the active site, formed by Cys102, 107 and His511, and a site at the dimer interface formed by His126 and Cys134 of both protein chains. PDB:3DK4. D GSR activity for wild-type human recombinant GSR upon titration with TPEN and ZnCl_2_ E GSR activity for wild-type and C134S mutant human recombinant GSR upon titration with ZnCl_2_ F Isothermal titration calorimetry (ITC) analysis for wild-type and H511K mutant human recombinant GSR G Energy minimized model of zinc binding to the active site of GSR, coordinated by Cys102, Cys107, Thr383, and His511. Based on PDB:2AAQ. H Treatment of SKMES1 and A549 lung cancer cells with zinc pyrithione shows a concentration dependent oxidation of the cellular glutathione pool after 5 hours of treatment I Treatment of SKMES1 and A549 lung cancer cells with zinc pyrithione or 2 µM pyrithione combined with ZnCl_2_ shows concentration dependent cytotoxicity during treatment for 24 hours. J Treatment of SKMES1 and A549 lung cancer cells with 2 µM pyrithione combined with ZnCl_2_ in presence of 10 mM N-acetyl cysteine (NAC) or 1 mM Glutathione ethyl ester (GSHee) limits zinc-mediated cytotoxicity K Overexpression of GSR in A549 lung cancer cells reduces zinc-mediated cytotoxicity upon treatment with 2 µM pyrithione combined with ZnCl_2_

Next, we aimed at leveraging zinc-mediated GSR inhibition to target GSR in lung cancer cells. For this we first determined the effect of zinc on the glutathione redox state in the GSR-dependent SKMES1 and GSR^High^ A549 lung cancer cell lines (**Figure S5C, D**), using the well-established GSH recycling assay^65–67^. We confirmed a significantly lower GSH pool size in A549 compared to SKMES1 cells, in line with publicly available metabolomics data^60^(**Figure S5D, S6A**). Upon 5-hour treatment with zinc pyrithione (a membrane permeable zinc complex), we observed a concentration-dependent depletion of free intracellular GSH compared to untreated samples, which correlated with an increase in GSSG levels in both SKMES1 and A549 cells (**Figure 7H**). As zinc effectively depleted free GSH in both cell lines, we hypothesized that zinc might induce substantial oxidative stress and thereby drive cytotoxicity GSR-reliant cancer cells. To examine this, we determined cell viability upon treatment with zinc pyrithione or pyrithione + ZnCl_2_ for 24 hours and found that both cell lines exhibited considerable sensitivity to zinc with LD_50_ at 10-20 µM, which coincided with almost complete depletion of total GSH (**Figure 7I, S6B**). Because pyrithione alone appeared slightly inhibitory to cell proliferation at elevated concentrations (**Figure S6C**), and due to the limited cellular penetrance of zinc salicylate and ZnCl_2_ (**Figure S6D, E**), a combination of low pyrithione + ZnCl_2_, was selected as standard treatment from here on. To examine a direct mechanistic link between zinc-mediated cytotoxicity and GSR inhibition, rescued zinc-mediated GSH depletion by replenishing cellular thiols using either N-acetyl cysteines (NAC; 10 mM) or cell permeable glutathione ethyl ester (GSHee; 1 mM), which both markedly alleviated zinc-induced toxicity (**Figure 7J, S6F)**. Moreover, a desensitization of A549 cells towards zinc was observed upon overexpression of GSR (**Figure 7K, S6G, H)**. In conclusion, leveraging ZnCPT together with pharmacological and genetic strategies we identify a novel mechanism of zinc-based inhibition of GSR that drives cytotoxicity in GSR-reliant lung cancer cells.

## Discussion

ZnCPT represents a comprehensive atlas of the zinc binding proteome, which defines thousands of constitutive and inducible zinc binding cysteines across a range of protein functional domains. Importantly, ZnCPT reports deep proteome coverage with high specificity and accuracy, enabling precise residue-level assignment of zinc binding to proteins. This compendium constitutes an extensive resource, which can guide future research into the understudied regulatory role of zinc over proteins and cellular physiology.

By capturing a substantial proportion of the proteome, ZnCPT defines general physicochemical and structural principles that govern zinc coordination. The systematic structural analysis of zinc binding site environments identified distinct features of zinc coordination sites between inducible and constitutive sites, providing mechanistic insights that can be further explored for individual protein and protein classes. These features also differentiate zinc binding from redox regulated cysteines, thereby enabling the classification of functionally distinct cysteine populations across the proteome.

The comprehensive mapping of the zinc-regulated proteome enables systematic investigation of zinc-regulated cellular processes. Founded on these analyses, ZnCPT unveils a link between zinc and cancer, identifying numerous cancer dependencies as targets of zinc regulation. Based on this, we elucidated the mechanistic basis for potent zinc-mediated inhibition of GSR by binding to its active site. This forms the basis for marked zinc-induced cytotoxicity in GSR-reliant lung cancer cells. In conclusion, further research into a potential therapeutic role for zinc to enhance chemotherapeutics-induced cytotoxicity by sensitizing cancer cells to redox stress will be of interest.

Taken together, the ZnCPT this compendium provides the basis for the mechanistic characterization of targets underlying zinc regulation and can be used as a foundation for future work elucidating the critical role of zinc over cellular functions in health and disease.

## Methods

### Resource Availability Lead contact

Further information and requests for resources should be directed to and will be fulfilled by the Lead Contact, Edward T. Chouchani.

### Materials availability

This study did not generate any new reagents.

### Experimental Model and Subject Details Maintenance of cell lines

HCT116 and Lenti-X™ 293T cells were cultured in DMEM (Corning, 10-017-CV) without pyruvate, supplemented with 10% FBS (GeminiBio, 100-106) and 1% P/S (Corning, 30-002-CI). SKMES1 and A549 cells were cultured in EMEM (ATCC, 30-2003), supplemented with 10% FBS (GeminiBio, 100-106) and 1% P/S (Corning, 30-002-CI). All cells were washed with PBS (Corning, 21-040-CV) detached using 0.25% trypsin (Gibco, 25200-056) and subcultured every other day.

### Method Details CPT synthesis

CPT was synthesized as described previously^10^. Briefly, 6-aminohexylphosphonic acid hydrochloride salt (6-AHP; SiKÉMIA) was added to succinimidyl iodoacetate (SIA; Combi-Blocks) to final concentrations of 45 mM SIA and 175 mM 6-AHP and reacted at room temperature for 1 hour while slowly stirring in the dark. The reaction was quenched with TFA (final pH < 2) and purified via HPLC (C18 column, solvent A: water with 0.035% TFA, solvent B: acetonitrile (ACN) with 0.035% TFA, 100%–40% solvent A over a 60-min gradient at a flow rate of 40 mL/min). The eluent containing CPT was frozen and lyophilized yielding a white powder. Quality was controlled via LC-MS as described previously^10^.

### Preparation of native cell lysates

HCT116 cells were cultured in standard medium (Dulbeco’s Modified Eagles Medium (DMEM) w/o sodium pyruvate) in 15 cm dishes to 80-90% confluency (5 dishes per experiment). Cells were washed with 10 ml ice cold PBS and then gently scraped into 5 ml of PBS. Cells were centrifuged for 5 min at 1000 x g followed by two washes with 50 mM HEPES, 150 mM NaCl (pH 7.5). The final cell pellet was resuspended in 0.8 ml of native lysis buffer (50 mM HEPES, 150 mM NaCl, 0.5 % (v/v) IGEPAL (pH 7.5), and 0.5 mM TCEP). Cells were incubated rotating for 10 min at 4°C and subsequently disrupted by passing 10x through a G28 injection needle. Cell lysates were clarified by centrifugation for 15 min (21.000 x g and 4 °C) and protein content in the supernatant was determined using a Pierce™ BCA assay kit (Thermo Scientific, USA). Cell lysates were diluted to a final concentration of 2 mg protein/ ml in native lysis buffer and kept on ice until used freshly.

### ZnCl_2_/chelator treatments and CPT labelling

Native cell lysates (200 µl corresponding to 400 µg protein) was distributed into individual Eppendorff tubes on ice. Zinc/Metal binding in native lysates was manipulated by addition of 100x stocks of ZnCl_2_ (10 µM final), TPEN (1mM final), EDTA (1 and 5 mM final), control samples received equivalent volume of buffer. Samples were incubated for 15 min at 37°C shaking at 500 rpm, prior to addition of 20 mM CPT (200 mM stock in 25 mM HEPES (pH 7.5), 150 mM NaCl, pH-ed with 1 M NaOH to pH 7.5. Accessible cysteines were labeled with CPT for 1 hour at room temperature in the dark. The labelling was rapidly quenched by precipitation of proteins on ice, using 3x vol. of methanol, 1 vol. of chloroform and 2.5 vol. of H_2_O, followed by vortexing and centrifugation for 15 min (15.000 x g and 4 °C). The resulting protein precipitate was washed three times with 1 ml methanol followed by centrifugation for each 5 min (15.000 x g and 4 °C).

### Protein digestion and TMT labelling

Precipitated protein pellets (∼400 µg protein) were dried and resuspended in 100 µl 200 mM N-(2-Hydroxyethyl)piperazine-N′-(3-propanesulfonic acid) (EPPS) (pH 8.0), containing trypsin (Promega; final 1/100 enzyme/protein ratio) and LysC (Wako, Japan; final 1/100 enzyme/protein ratio). Protein lysates were digested overnight shaking vigorously (1500 rpm at 37 °C). Samples were centrifuged for 10 min at 21,000 x g, soluble peptide suspension was transferred into fresh Eppendorf tubes and peptide concentration was determined by microBCA (Thermo Scientific). Equal amounts of peptides for each sample (150-200 µg) were transferred into fresh tubes and samples were completed to a volume of 100 µl with 200 mM EPPS (pH 8.0). Peptides were labelled with 40 µl of TMTpro (16- or 18-plex) reagents in acetonitrile for 1 hr in the dark at RT, while vortexed intermittently. Following the labelling 2 µl of each sample were combined into 120 µl of 1% formic acid in H_2_O, desalted and analysed as ratio check via LC-MS. TMT labelled samples were stored transiently at −80 °C. Upon completion of the ratio check analysis, the labelling reaction was quenched by addition of 5 µl hydroxylamine (5% stock) and incubation for 15 min at RT. Samples were pooled at equal amounts according to the ratio check and diluted with 12 ml of 1% formic acid in H_2_O and subjected to gravity flow driven C18 solid-phase extraction (200 mg Sep-Pak, Waters) and subsequently vacuum dried.

### Cysteine peptide enrichment

TMT labelled pooled peptides were resuspended in phosphatase buffer (50 mM HEPES, 100 mM NaCl, 1 mM MnCl_2_ (pH 7.5) and Lambda phosphatase (Santa Cruz Biotechnologies) was added according to manufacturer’s instructions. Peptides were dephosphorylated during 2 hrs at 30 °C shaking at 500 rpm. Subsequently, the sample was acidified with 10% TFA to a pH of < 3.0 (∼60 µl), subjected to gravity flow driven C18 solid-phase extraction (200 mg Sep-Pak, Waters) and vacuum dried. CPT labelled cysteine peptides were purified using the High-select Fe-NTA phosphopeptide enrichment kit (Thermo Scientific) according to manufacturer’s instructions. Following elution of enriched CPT-labelled cysteine peptides, the peptide suspension is acidified with 10% TFA to a pH of < 3.0 (∼25 µl), subjected to gravity flow driven C18 solid-phase extraction (50 mg Sep-Pak, Waters) and vacuum dried.

### Cysteine peptide fractionation by HPLC

Dried peptides (∼100 µg) were resuspended in 300 µl of high-performance liquid chromatography (HPLC) buffer A containing 5 mM ammonium bicarbonate pH 8.0, 5% acetonitrile and centrifuged through a PTFE 0.2 µM filter (Merck). Peptides were fractionated with basic pH reversed-phase HPLC using an Agilent 300 extend C18 column. A 50-min linear gradient in 13 - 43% buffer B (5 mM ammonium bicarbonate, 90% acetonitrile, pH 8.0) at a flow rate of 0.25 ml/min, and eluates were collected into a 96-deep-well plate. Fractions were consolidated into 12 tubes and vacuum dried followed by peptide desalting (stage tip) and LC-MS/MS analysis.

### Peptide desalting (stage tip) for LC-MS/MS

Peptides were desalted prior to LC-MS analysis using solid phase extraction (stage tip). Briefly, C18 Octadecyl HD solid phase extraction disk (CDS Analytics, USA) was used to prepare stage tips in house. The matrix was activated with 100% ACN, followed by washes with 70% ACN, 1% FA and H_2_O, 1 % FA. Dried peptides were dissolved in H_2_O, 1 % FA and passed through the C18 matrix. Peptides were washed twice with H_2_O, 1 % FA and subsequently eluted into MS vials in two steps with 40% ACN, 1% FA and 70% ACN, 1% FA. Peptides were vacuum tried and stored at −80°C until analysis.

### LC-MS/MS parameters

An Orbitrap Eclipse Tribrid Mass Spectrometer (Thermo) coupled with an Easy-nLC 1200 (Thermo) was used for proteomics measurements. Of each fraction ∼ 3 µg of peptides, dissolved in 5% ACN, 5% FA were loaded onto an in-house 100-μm capillary column packed with 35 cm of Accucore 150 resin (2.6 μm,150 A::J). Peptides were separated and analyzed using a 180-min gradient consisting of 2% - 23% ACN, 0.125% FA at 500 nl/min flow rate. A FAIMSPro (Thermo) device was used for field asymmetric waveform ion mobility spectrometry (FAIMS) separation of precursors^70^, and the device was operated with default settings and multiple compensation voltages (−40V/-60V/-80V). Under each voltage, data-dependent acquisition mode was used for a mass range of m/z 400-1400 applying a top10 DDA method. Resolution for MS1 was set at 120,000. Singly-charged ions were not further sequenced, and multiply-charged ions were selected and subjected to fragmentation with standard automatic gain control (AGC) and 35% normalized collisional energy (NCE) for MS2, with a dynamic exclusion window of 30 s. Quantification of TMT reporter ions were performed using the multinotch SPS-MS3 method with 45% NCE for MS3, which is optimized for TMTpro-16/-18 reagents.

### Database searching

Raw files were first converted to mzXML, and searched using the Comet algorithm^72^ on an in-house database search engine reported previously^73^. Database searching included all human (Homo Sapiens) entries from UniProt (http://www.uniprot.org, downloaded 2021) and the reversed sequences as well as common contaminants (keratins, trypsin, etc.). Peptides were searched using the following parameters: 25 ppm precursor mass tolerance; 1.0 Da product ion mass tolerance; fully tryptic digestion; up to three missed cleavages; variable modification: oxidation of methionine (+15.9949); static modifications: TMTpro (+304.2071) on lysine and peptide N terminus. The false discovery rate (FDR) was controlled as described previously^73–75^ to < 1% on peptide level for each MS run using parameters such as XCorr, ΔCn, missed cleavages, peptide length, charge state and precursor mass accuracy. Then protein-level FDR was also controlled to < 1%. Cysteine site localization was determined using the ModScore algorithm^76^ where a score of 19 corresponds to 99% confidence in correct localization.

### TMT reporter-based quantification

TMT reporter ions were used for quantification of peptide abundance. Each reporter ion was scanned using a 0.003 Da window, and the most intense m/z was used. Isotopic impurities were corrected according to the manufacturer’s specifications, and signal-to-noise ratio (S/N) was calculated. Peptides with summed S/N lower than 160 across 16 channels of each TMTpro16 plex (180 across 18 channels of each TMTpro18 plex) or isolation specificity lower than 0.5 were discarded.

### Data analysis

All data analyses were performed in R (Version 4.2.1) unless stated otherwise. Spearman correlations were either performed in GraphPad Prism 10 or in R. Significance of cysteine exposure was determined with two-tailed Student’s t tests for pairwise comparison, and multiple comparisons were corrected using the Benjamini-Hochberg procedure^77^. Sites with accessibility changes >1.5 (constitutive zinc/metal binding: TPEN/EDTA) or <0.8 (inducible zinc binding: ZnCl_2_) and p.adj < 0.05 were defined as significantly changed sites. <u>The heatmap</u> highlighting accessibility changes was generated for cysteine sites quantified across the entire dataset (no missing values). For the clustering the pheatmap package (Version 1.0.12) using the default complete linkage clustering method. Uniform Manifold Approximation and Projection (UMAP) analysis for dimension reduction was performed for cysteine sites quantified across the entire dataset (no missing values) using the UMAP package^78^. <u>Protein subcellular localization</u> data was downloaded from Human Proteome Atlas (https://www.proteinatlas.org) based on data previously reported^20^ and matched to the ZnCPT dataset. Nucleoplasm, nuclear speckles, nuclear bodies, nuclear membrane, nucleus, nucleoli fibrillar center, and nucleoli rim were consolidated into the nucleus annotation. Actin filaments, centrosome, cytosol, cytoplasmic bodies, cytokinetic bridge, centriolar satellite, intermediate filaments, midbody, microtubules, microtubule ends, midbody ring, mitotic chromosome, microtubules ends, mitotic spindle, and rods & rings were consolidated into the cytoplasm annotation. Golgi apparatus and Golgi were consolidated into the Golgi annotation. Endosomes and lysosomes were consolidated into the EndoLysosomes annotation. Cell junctions, focal adhesion sites, and plasma membrane were consolidated into the plasma membrane annotation. Distribution of proteins containing significantly or non-significantly changing cysteines across subcellular compartments was assessed, and enrichment of proteins for the individual compartments was calculated using Fisher’s exact tests. For <u>Pfam protein domain</u> analysis, the Pfam dataset for the human proteome (Taxonomy ID: 9606) was retrieved from the InterPro website (https://www.ebi.ac.uk/interpro). Pfam domains were matched to the ZnCPT dataset based on Uniprot ID and the position of quantified cysteines and the start/end of the domain within the protein sequence. Pfam domains matching at least three cysteines within ZnCPT were selected and Pfam enrichment for significantly changed cysteines was calculated using Fisher’s exact tests. Multiple comparisons were corrected using the Benjamini-Hochberg procedure^77^. <u>GO Process and GO Function</u> term datasets for the human proteome (Taxonomy ID: 9606) were obtained from the QuickGo website (https://www.ebi.ac.uk/QuickGO/annotations). GO terms matching at least three proteins within ZnCPT were selected and GO term enrichment for proteins containing significantly changed cysteines was calculated using Fisher’s exact tests. Multiple comparisons were corrected using the Benjamini-Hochberg procedure^77^. The ZincBind^6^ dataset was kindly provided by Sam Ireland and Andrew Martin. All human Uniprot data was retrieved from Uniprot (https://www.uniprot.org). Uniprot Name, Uniprot ID and all associated PDB information was extracted. ZincBind was filtered for PDB IDs that were contained within the human Uniprot dataset and in which zinc was coordinated by a cysteine residue. The ZincBind and human Uniprot datasets were merged by PDB ID and chain ID of the metal binding residue defined in ZincBind. The generated final ZincBind dataset was matched to the ZnCPT dataset by Uniprot ID and position of quantified cysteines. The overlap of both datasets for cysteine sites and proteins, was calculated for the entire dataset and sites or proteins containing sites that exhibit significantly changed accessibility of any magnitude. <u>Zinc binding</u> <u>motifs</u> (+/- 6 amino acids around the quantified cysteine) were extracted and separated into significantly changing and background motifs. Comprehensive motif analysis was perform using the MEME suite (https://meme-suite.org/meme/) and the XSTREME^34^ motif analysis algorithm. Sequence logos of significantly enriched motifs were retrieved. <u>The oximouse dataset</u>^10^ was obtained from the oximouse website (https://oximouse.hms.harvard.edu). To match the oximouse dataset to ZnCPT, mouse and human entire proteome sequences were retrieved from Uniprot (https://www.uniprot.org). Human and mouse proteins were matched by their Uniprot names, all selenocysteines (U) were replaces with cysteines (C), and sequences were aligned using the Biostrings package. The site number of conserved cysteines was extracted and used to match cysteine oxidation values from the oximouse dataset to ZnCPT. Delta oxidation values for cysteines were calculated for young and aged mice across all organs to map the redox regulatory potential for each cysteine. For <u>KEGG</u> <u>pathway</u> enrichment analysis, the human KEGG pathway dataset mapped to KEGG gene identifiers was retrieved from the KEGG website (https://rest.kegg.jp/). The KEGG pathways were mapped to a dataset containing Uniprot IDs and Uniprot names, by KEGG gene identifiers retrieved from Uniprot (https://www.uniprot.org). From this, KEGG pathway annotations were mapped to ZnCPT. KEGG pathways matching at least three proteins within ZnCPT were selected and KEGG pathway enrichment for proteins with significantly changed cysteines was calculated using Fisher’s exact tests. Multiple comparisons were corrected using the Benjamini-Hochberg procedure^77^. The <u>DepMap chronos cancer dependency</u> dataset was retrieved from the DepMap portal ((https://depmap.org/portal) Public 13Q2 release: CRISPR (DepMap Public 23Q2+Score, Chronos dataset). Z-scores were calculated for each gene across all cell lines within the datasets. The DepMap dataset was mapped to ZnCPT by gene name. The Gene Expression dataset was retrieved from the DepMap portal ((https://depmap.org/portal) Public 13Q2 release: Expression Public 23Q2. The <u>cancer cell line proteome</u> dataset^59^ was obtained from the Cell Model Passports website (https://cellmodelpassports.sanger.ac.uk; release:2022.12.14). The <u>cancer cell line metabolome</u> dataset^60^ was obtained from the DepMap portal (https://depmap.org/portal) Public 13Q2 release: Metabolomics dataset). Intensities for abundance of GSH, GSSG, and NADP were extracted for all cancer cell lines within the dataset. Protein structural data was retrieved from the RCSB protein databank (https://www.rcsb.org) or AlphaFold^69,79^ (https://alphafold.ebi.ac.uk). The AlphaFold structure for MTF1 (AF-Q14872-F1) was processed with the AlphaFill^33^ web application (https://alphafill.eu) to model zinc by similarity into the zinc finger domains predicted by AlphaFold. Structural analysis including figure generation was performed in PyMOL (Version 2.3.1).

### Analysis of the 3D structural coordination motif of zinc binding

Structural motif analysis were performed in Python (Version 3.7). The AlphaFold structures for human proteins were downloaded from alphafold.ebi.ac.uk. Each structure was cleaned by pruning the low confidence regions (pLDDT < 70). Cysteine sites identified by ZnCPT were mapped to the AF structures, and residues within 5 Å of the sulfur atom of the cysteine were extracted as the microenvironment for the cysteine of interest. The environment matrix for each cysteine was calculated by binning the occurrence of each amino acid type between 2 Å and 5 Å with a step of 0.5 Å. The proximity of each residue was measured by the shortest distance between the sulfur atom and the residue heavy atoms. A combined matrix for all constitutive and inducible sites was generated. To cluster the structure environment, the sequence of amino acid was constructed based on the environmental matrix. The sequence was ordered in proximity shells to the cysteine (i.e. sulfur atom). The first shell was between 2 – 3 Å, the second shell was between 3 – 4 Å, and the final shell is between 4 – 5Å. A special character “X” was inserted between each shell to indicate shell structure. For example, the sequence “CCXFVXDW” indicated a disulfide bond in the first shell because of short distance between two CC, and two residues phenylalanine (F) and valine (V) in the second shell, and aspartate (D) and tryptophan (W) in the third shell. The order of amino acid within each shell followed a predefined order independent of primary sequence. The sequences was then aligned using biopython globalxx algorithm without gap penalty. The alignment scores were converted to distance in terms of percentage of match residues, and the distance matrix was used to cluster the sites. The clustering was done using scipy cluster package with “ward” algorithm. Uniform manifold approximation and projection (UMAP) plots were generated using python umap package to project the distance matrix onto 2D.

### Molecular modeling GSR zinc binding site

GSR crystal structure 2AAQ was used as the template for generating the molecular model initial structure, where Au ion was replaced by Zn^2+^. The nearby residues (residue 467 of chain A, residue 339, 58, 63 of chain B) were chosen at quantum mechanical region. Other residues were treated classically. Energy optimization was carried out using B3LYP density functional model with LACVP++** basis set by QM/MM module in the Schrödinger PyMOL software suite (Version 2.5).

### *In vitro* hydroxylation of HIF1**α**-ODD peptide by human recombinant EGLN1

To measure prolyl hydroxylation activity, recombinant human PHD2(EGLN1) was purchased from Active Motif (USA, #81065) and HIF1α-ODD peptide (DLDLEALAPYIPADDDFQL) was purchased from GeneScript Biotech (USA)). A mastermix reaction solution containing 5 µg/ml recombinant EGLN1, 20 µg/ml HIF1α-ODD peptide, 5 mM KCl, 1.5 mM MgCl_2_, 100 µM 2-Oxoglutarate, 100 µM L-Ascorbate and 50 µM Ammonium iron(II) sulfate hexahydrate containing additionally indicated concentrations of ZnCl_2_ was prepared. The reaction was started upon addition of 2-Oxoglutarate. At indicated time points 10 µl of the reaction solution were retrieved, quenched by addition of 5% ACN 5% FA and stored at −20°C until same-day analysis via LC-MS. Samples were separated on a a PLRP-S 1000A, 2.1[×[50[mm, 5 μm column (Agilent).The mobile phases were MS solvent A (H_2_O, 2% FA) and B (ACN, 2 % FA) at a flow rate of 0.3 ml/min with the following gradient (the proportion of MS solvent B is given in %): 0–1.5 min: 15%, 1.5–3 min: 15–95%, 3–3.5 min: 95%, 3.5–4 min: 100-15%, 4–5 min: 15% at 60°C column temperature. For analysis a Q-Exactive HF-X mass spectrometer (Thermo Fisher Scientific) in positive ion mode was used with full scan analysis over a range of m/z[400– 1600[m/z at 60,000 resolution, 1[×[10^5^ AGC target and 50 ms maximum ion accumulation time. Top 5 multiply charged peptide ions were selected for MS/MS each second and were analyzed using the following parameters: resolution 15,000; AGC target of 1[×[10^5^; maximum ion transfer of 100 ms; 0.7 m/z isolation window; for HCD a normalized collision energy 34% was used; and dynamic exclusion of 10 s. HIF1α-ODD peptides were quantified by determining the peak area of XICs (extracted ion chromatograms) of monoisotopic peaks (1067.5149 m/z HIF1α-ODD peptide, 1075.5124 m/z Hydroxy-HIF1α-ODD peptide; 10 ppm mass error) using the Themo Xcalibur software (v4.1)

### Expression and purification of human recombinant GSR

The N-terminal 6xHis-tagged construct of human GSR (wild-type and C134S mutant) (residues 44–522) was cloned into a pET-28a(+) expression vector was purchased from GeneScript Biotech (USA). The N-terminal TwinStrep-tagged construct of human GSR (wild-type and H511K mutant) (residues 44–522) was cloned into a pET-28a(+) expression vector was purchased from GeneScript Biotech (USA). Proteins were overexpressed in E. coli BL21(DE3)(Life Technologies, USA) and purified using affinity chromatography. Cells were grown at 37°C in TB medium (containing 50 µg/ml kanamycin) to an OD600 of 1, cooled to 16°C and induced with 1 mM isopropyl-1-thio-D-galactopyranoside (IPTG) and grown over night. Alternatively, cells were grown in Overnight Express™ Auto-inducible medium (containing 50 µg/ml kanamycin) to an OD600 of 1, cooled to 16°C and grown over night. Cells were harvested by centrifugation (6000 x*g*) and resuspended in ice-cold lysis buffer (100 mM HEPES, 500 mM NaCl, 5 mM TCEP (and 20 mM Imidazole for 6xHis-taged proteins), pH 8. Cells were lysed by sonication using a Q500 Sonicator (QSONICA Sonicators, USA) during 10x 30s on, 30s off cycles. The resulting lysate was centrifuged for 30 min (30,000 x g, 4°C) and the soluble fraction was collected. <u>His-tagged:</u> The lysate was mixed head-over-head with Ni-NTA agarose resin (Thermo Scientific™, USA) for 60 min at 4°C. Resin was transferred to chromatography columns and washed with 15 column volumes of 100 mM HEPES, 500 mM NaCl, 5 mM TCEP, and 20 mM Imidazole, pH 8. Protein was eluted in 10 fractions with each 1.5 ml of 100 mM HEPES, 100 mM NaCl, 5 mM TCEP, and 250 mM Imidazole, pH 8. <u>TwinStrep-tagged:</u> The lysate was passed through chromatography columns containing 1.5 ml bed volume of Strep-Tactin® Sepharose® resin (IBA Lifesciences, Germany). Resin was washed with ∼20 resin volumes of 100 mM HEPES, 500 mM NaCl, 1 mM TCEP, pH 8 and eluted in 7 fractions of 0.5 ml of 100 mM HEPES, 150 mM NaCl, 2.5 mM Desthiobiotin, 1 mM TCEP, pH 7.5. <u>All purified protein:</u> Fractions containing glutathione reductase were combined and desalted using 5 or 10 ml Zeba™ Spin desalting columns (Thermo Scientific™, USA) into 50 mM HEPES, 150 mM NaCl, 5 mM TCEP, pH 7.4. TwinStrep-tagged proteins were concentrated using 30K MWCO concentrators to ∼ 5 mg/ml (Thermo Scientific™, USA). Protein was stored at −80°C.

### Isothermal Titration Calorimetry

For ITC, protein samples were filtered (0.2 µM centrifugal filter) and were then further purified by size-exclusion chromatography on Superdex 200 resin (GE Healthcare) using 25 mM HEPES, 150 mM NaCl, pH 7.5 as buffer. ZnCl_2_ (Sigma) was dissolved in the same buffer. ITC experiments were carried out in an Affinity ITC instrument (TA Instruments) at 25[°C. The titrations were performed by injecting 2.5 μl aliquots of 10x (concentration of protein) ZnCl_2_ into the calorimeter cell containing a 185 μl solution of 40.38 µM wild-type or 29.74 µM H511K glutathione reductase with a constant stirring speed at 125 rpm, and the heats were recorded. The data were analyzed with the NanoAnalyze using the independent fit model. All the uncertainties were estimated by the native statistics module with 10000 synthetic trials and 95% confidence level.

### Cell Viability Assay

Cell viability was assessed using the CellTiter-Glo® Luminescent Cell Viability Assay (Promega). Cells were seeded at a density of 7000 cells/ well (in 50 µl standard culture medium) into white 384-well flat bottom, low flange plates (Corning) and left to attach for at least 6 hours. The medium was exchanged for 25 µl standard medium containing indicated concentrations of chemicals (6-plicate per condition) for up to 24 hours. Cell viability was determined upon addition of 25 µl CellTiter-Glo® reagent in accordance with manufacturer’s instructions and luminescence was measured using a ClarioSTAR Plus plate reader (BMG Labtech, Germany).

### Glutathione Reductase Activity Assay

Glutathione Reductase activity (recombinant human Glutathione Reductase (Bio-Techne: #8866-GR-100) or homemade recombinant Glutathione Reductase) was determined by measuring the consumption of NADPH in the presence of GSSG. The assay was performed in a 96-well format with a total assay volume of 200 µl per well and three technical replicates of each condition. The NADPH absorbance was measured with a ClarioSTAR Plus microplate reader (BMG Labtech, Germany). All reagents were prepared in 100 mM Tris-HCl pH 7.5. Wells were pre-plated with 20 µl of 10x ZnCl_2_ or TPEN dilutions, followed by addition of 30 µl 100 mM Tris-HCl pH 7.5. Then, 50 µl of recombinant Glutathione Reductase at a concentration of 1.2 µg/ml and 50 µl of 4 mM GSSG was added. The absorbance was measured for 50 cycles at room temperature monitoring the absorbance at λ = 340 and 380 nm in 45 second intervals. Following 5 cycles of baseline readings, 50 µl of 0.8 mM NADPH was rapidly added to each well and measurements were continued. The maximum linear rate of NADPH oxidation and the ΔAbsorbance (340–380nm) was calculated. The NADPH concentration was determined using the extinction coefficient ε_340-380_ = 4.81 mM^-1^cm^-1^.

### Measurement of total GSH and GSSG in cells

The total (GSH) and oxidized (GSSG) glutathione of cells was determined using the well-established glutathione recycling assay. Cells were seeded into 6-well plates at 5×10^5^ cell/well the day before the assay (2.5×10^5^ cell/well two days before the assay (24-hour treatment)). Cells were treated for 5 or 24 hours with indicated concentrations of zinc pyrithione, pyrithione + ZnCl_2_. Following the treatment, cells were rapidly collected and lysed in 170 µl ice-cold 5 % 5-sulfosiacylic acid (SSA) and stored on ice. The 6-well plates were processed on a plate-by-plate basis to ensure rapid quenching of the glutathione redox state. Upon completion of cell lysis, extracts were cleared by centrifugation for 10 min (21.000 x g and 4 °C) and supernatants were transferred into fresh Eppendorf tubes on ice. Protein pellets were soaked in 10 µl of 20% SDS for 5 min and completed to 100 µl with 100 mM Tris-HCl pH 7.5 and proteins were dissolved by shaking (1500 rpm) at room temperature for 1 hour. The protein concentration was determined using a Pierce™ BCA assay kit (Thermo Scientific™, USA). <u>Total GSH levels:</u> Clarified cell extracts were diluted 1:3 in 5% SSA. GSH standards (0, 10, 20, 30, 40, 50, 60, 70 µM) were prepared in 5 % SSA. Extracts and diluted samples were plated at 10 µl/well into 96-well plates in duplicate. <u>GSSG levels:</u> GSH standards (0, 0.5, 1, 2, 4, 8, 10, 20 µM) were prepared in 5 % SSA. Reduced glutathione (GSH) was derivatized to only measure oxidized glutathione (GSSG) levels. For this, 60 µl of standards and non-diluted extracts were transferred into fresh Eppendorf tubes. Samples were neutralized and derivatized by addition of 10 µl derivatization mix (2.25 µl 2-vinyl pyridine + 2.75 µl of Triethanolamine +5 µl H_2_O) and incubated for 1 hour rotating head-over-head in the dark at 4 °C. Then, samples were plated at 10 µl/well into 96-well plates in duplicate. <u>Assay:</u> Wells were completed with 240 µl reaction buffer (all in 150 mM HEPES, 1 mM EDTA pH 7.5: 150 µl 0.4 mM NADPH, 40 µl buffer, 50 µl 3 mM DTNB). The absorbance was measured for 35 cycles at room temperature monitoring the absorbance of TNB^2-^ at λ = 412 nm in 35 second intervals. Following 3 cycles of baseline readings, 50 µl of 0.8 mM NADPH in 150 mM HEPES, 1 mM EDTA pH 7.5 was rapidly added to each well and measurements were continued. The linear rate of TNB^2-^ formation was calculated for standards and GSH[GSH_Total_] or GSSG[GSSG] levels of samples were interpolated from the standard curves. Glutathione level calculations: [GSH_Free_] = [GSH_Total_] – 2x[GSSG] | GSSG content (%) = [GSH_Free_]/ 2x[GSSG]*100.

### Overexpression of Glutathione Reductase in A549 cells

Glutathione reductase lentiviral expression vector (pLV[Exp]-EGFP:T2A:Puro-EF1A>hGSR[NM_000637.5]) was purchased from VectorBuilder Inc. (China). Virus was produced by Lenti-X™ 293T cells (Takara Bio USA), transfected with lentiviral expression vector (330 ng), as well as pPAx (550 ng) and pMD2 (330 ng) constructs using PolyFect™ transfer reagent (Qiagen). For transduction, viral supernatant was passed through a 0.45 µM syringe filter, Polybrene (10 µg/ml; Sigma) was added, and was transferred onto A549 cells seeded into a 24-well plate (1-2 ml/well). This step was repeated the following day. Cells were selected by addition of 2 µg/ml Puromycin (Gibco) for 10 days, changing the medium daily. Following selection, medium was replaced with standard culturing medium, a polyclonal fraction of cells was collected. Remaining cells were diluted to a concentration of 0.5 cells/100 µl and 100 µl were plated into 96-well plates to obtain single cell clones. Glutathione reductase overexpression was confirmed by western blot.

### Western blot

For western blot analysis, cultured cells were washed with PBS and lysed in ice-cold RIPA buffer supplemented with EDTA-free cOMPLETE protease inhibitor (Roche) on ice. Lysate was collected and clarified by centrifugation for 10 min (21.000 x g and 4 °C). The protein concentration was determined using a Pierce™ BCA assay kit (Thermo Scientific, USA). Lysate was adjusted to equal protein concentration across samples and diluted with 4x NuPAGE LDS sample buffer (Thermo Scientific, USA) containing 50 mM DTT (Sigma, USA) and samples were heated for 15 min at 65°C. Samples were separated on 4-12 % NuPAGE BisTris (Thermo Scientific, USA) gels using MOPS SDS running buffer (Thermo Scientific, USA). Proteins were transferred to PVDF membranes using the iBLOT2 transfer system (Thermo Scientific, USA) with iBLOT2 PVDF transfer stacks (Thermo Scientific, USA). Membranes were blocked with 3% BSA (Sigma, USA) in TBS + 0.1% Tween (Boston BioProducts, USA). Primary antibodies (GSR Polyclonal Antibody (Rabbit, Proteintech: #18257-1-AP); GAPDH Monoclonal Antibody (Mouse, Proteintech: #60004-1-Ig)) were diluted 1:1000 in TBS + 0.1% Tween (Boston BioProducts, USA) containing 3% BSA (Sigma, USA), and membranes were incubated with antibodies over-night at 4°C. Membranes were washed three times with TBS + 0.1% Tween (Boston BioProducts, USA) followed by incubation for 1 hour at room temperature in the dark in with secondary antibodies (Anti-rabbit: IgG DyLight 800, Anti-mouse: IgG DyLight 680 (Cell Signaling Technologies, USA)), dissolved at 1:15,000 dilution TBS + 0.1% Tween (Boston BioProducts, USA) containing 3% BSA (Sigma, USA). Membranes were washed three times with TBS + 0.1% Tween (Boston BioProducts, USA) and membranes were scanned using an Odyssey DLx (LI-COR Biosciences, USA) scanner.

### Quantification and Statistical Analysis

Data processing and statistical analysis was performed using the pipeline described in the section above. Alternatively, data was visualized, and statistical analysis was performed using the Prism 10.0 software (Graphpad, USA). All data are represented as mean ± S.E.M., unless specified otherwise. Significance was calculated using two-tailed Student’s t test for pairwise comparison of variables. Fisher’s exact test was used for enrichment analyses. P values of hypergeometric tests were corrected for multiple comparisons using the Benjamini-Hochberg procedure^77^. For proteomics analysis at least triplicates were analyzed for each condition. Within plate-reader based assays, technical replicates (duplicates or triplicates) of the same sample were analyzed. The p (associated probability) value was considered significant if < 0.05 and significance was indicated as follows: *p < 0.05; **p < 0.01; ***p < 0.001, ****p < 0.0001.

### Figures

Figures were prepared in Adobe Illustrator 2022, Graphpad Prism 10.0, PyMOL (Version 2.3.1 and Version 2.5), R (Version 4.2) and Python (Version 3.7).

## Acknowledgements

E.T.C: This work was supported by the Claudia Adams Barr Program, the Lavine Family Fund, the Pew Charitable Trust, NIH DK123095, NIH AG071966, The Smith Family Foundation, and the American Federation for Aging Research.

N.B. is supported by the Deutsche Forschungsgemeinschaft (DFG, German research foundation: Projektnummer 501493132).

M.J.M. is supported by the Deutsche Forschungsgemeinschaft (DFG, German research foundation: Projektnummer 461079553).

H.X. is supported by the NIH (NIH K99AG073461)

L.H.M.B. was supported by the American Heart Association (926512)

H-G.S. is supported by Hope Funds for Cancer Research HFCR-20-03-01-02.

Y.S. is supported by the NIH (K01DK132455)

N.D. was supported by the NIH (T32CA236754)

P.L.-M. is supported by the NIH NIDDK (K99DK133502)

J.J.P. is supported by the NIH (K00AG073493)

We thank Sam Ireland and Andrew C.R. Martin for providing the ZincBind dataset.

We thank Jan Lj. Miljkovic for comments and suggestions.

## Author Contributions

N.B. and E.T.C. carried out study conception and design.

N.B. designed and performed most experiments, analyzed data, interpreted results and prepared figures.

M.J.M. assisted with GSR validation experiments.

H.X. assisted with LC-MS experiments, data interpretation and analysis.

L.H.M.B. assisted with the generation of over-expression cell lines.

S.S. assisted with LC-MS experiments and cell viability assays.

S.W. assisted with data analysis.

H-G.S. assisted with GSR validation experiments.

Y.S. performed ITC measurements with assistance of N.B.

N.D. assisted with LC-MS experiments.

J.J.P. assisted with GSR validation experiments.

P.L.-M. assisted with recombinant protein work.

Y.Z. and J.C. performed structural motif analysis, modeling of zinc binding and interpretation. The manuscript was written by E.T.C and N.B. with help of all authors.

E.T.C. directed the study.

---

**Supplementary Figure 1:**
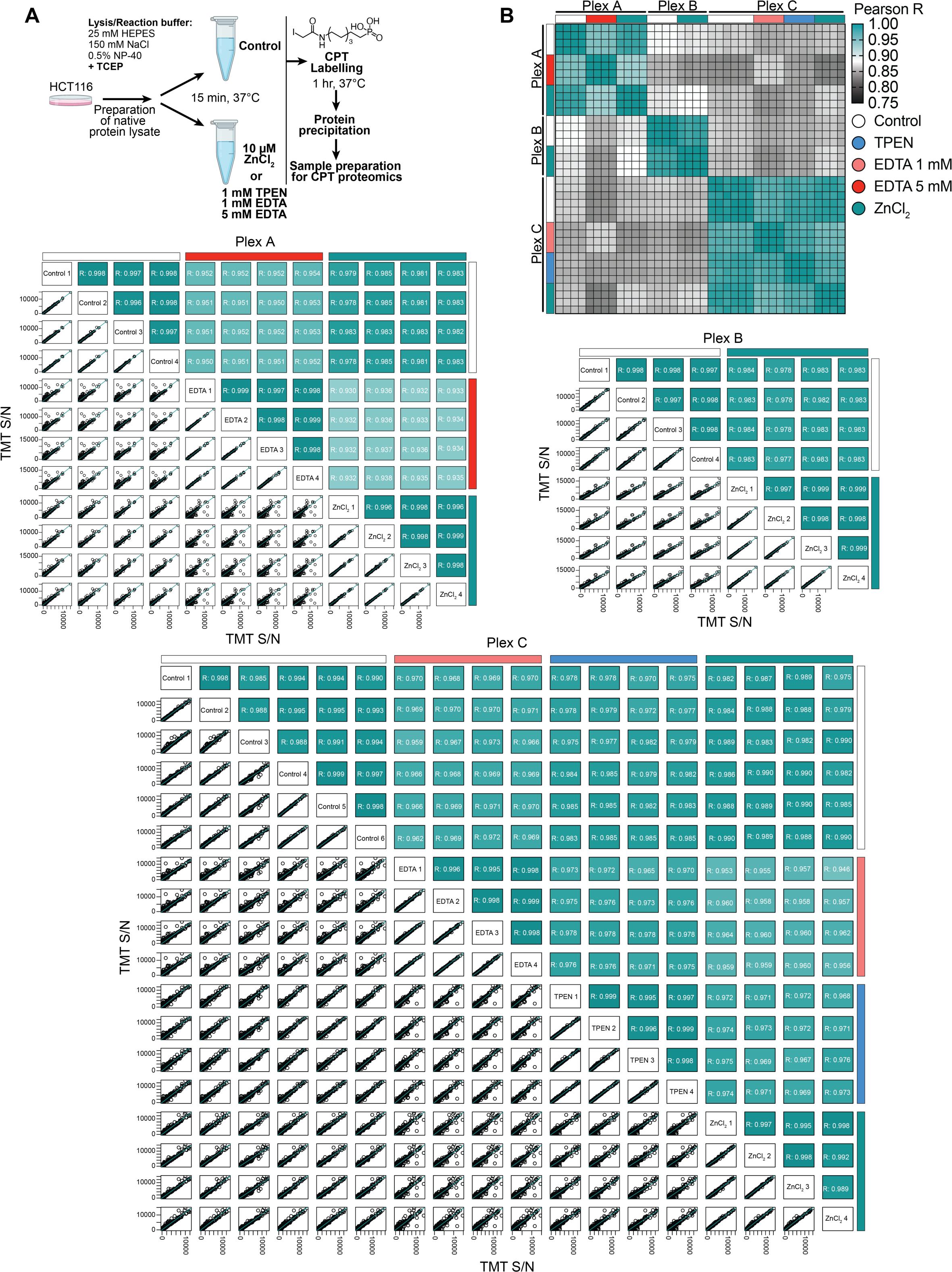
Experimental strategy and characterization of the ZnCPT dataset. A Experimental workflow for proteomic sample preparation to determine zinc coordination by protein cysteines B Quantification of cysteine containing peptides across different treatments in three plexes. Pairwise comparison demonstrates high reproducibility.

**Supplementary Figure 2:**
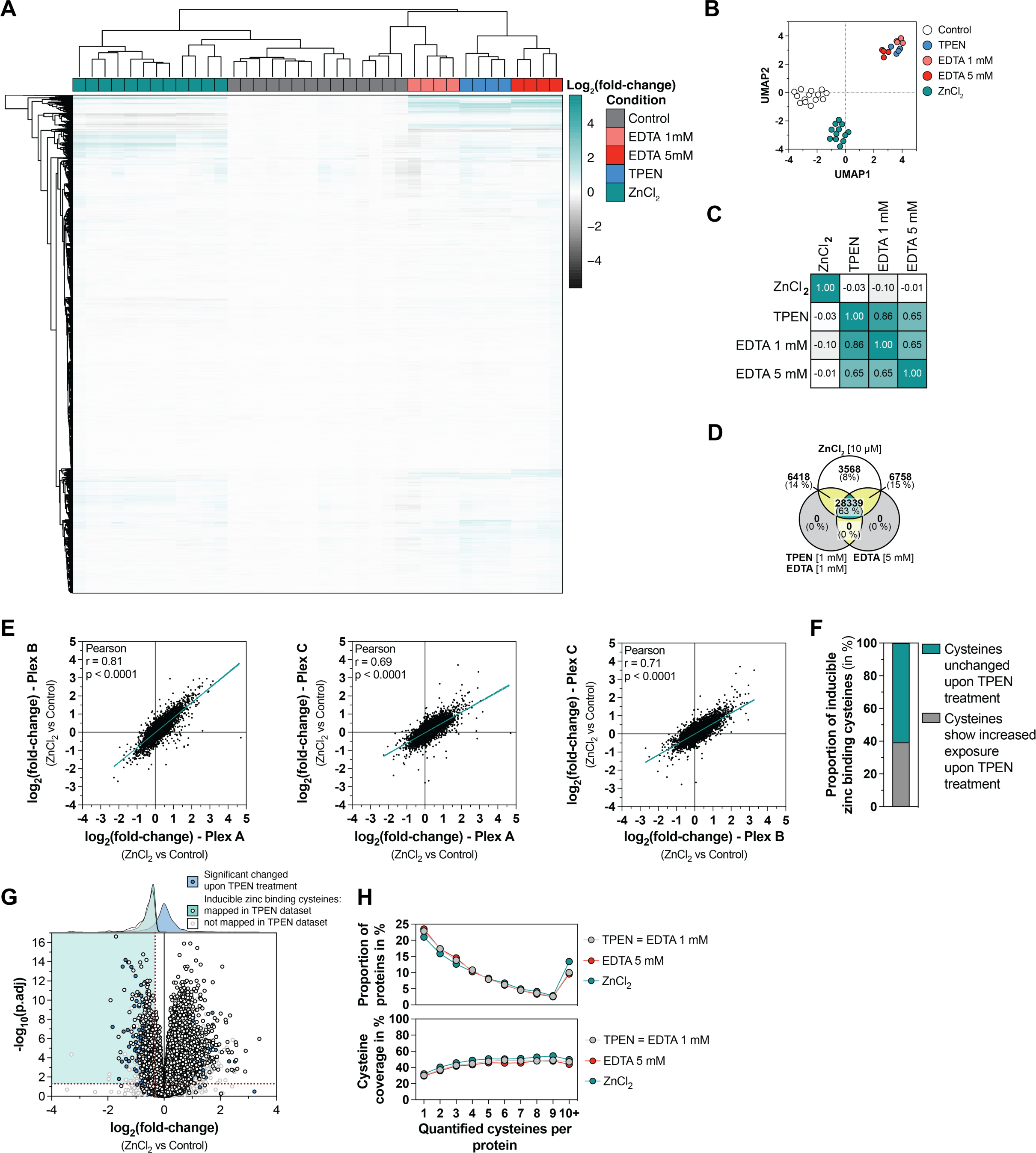
Characterization and benchmarking of the ZnCPT dataset. A Hierarchical clustering of cysteine site accessibility, normalized to the average of respective controls across all replicate samples, shows a distinct clustering of treatment conditions. B UMAP analysis across all replicates demonstrates a distinct clustering of treatment conditions, with all chelator treatments clustering together. C Correlative map of different treatment conditions, specifying Pearson coefficients. D Overlap of quantified cysteine sites across treatment conditions/plexes E Correlations of cysteine accessibility changes upon ZnCl_2_ treatment across three independent experiments F Classification of inducible zinc binding cysteines according to their accessibility changes upon TPEN treatment G Cysteine accessibility changes upon ZnCl_2_ cross referenced to their accessibility upon TPEN treatment reveals that many inducible zinc binding cysteines do not bind zinc under baseline conditions H Distribution of quantified cysteines per protein and cysteine coverage across treatment conditions

**Supplementary Figure 3:**
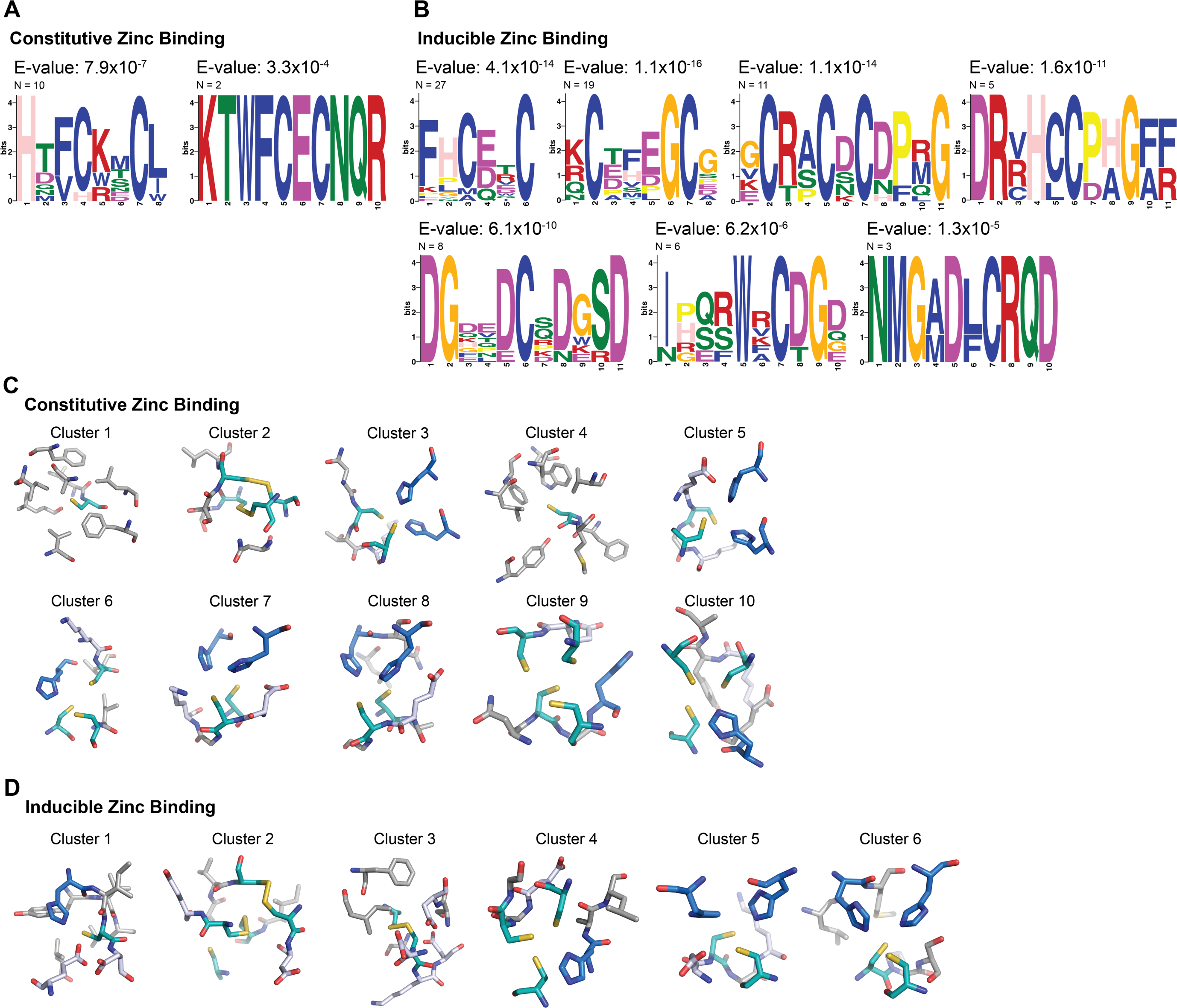
Primary sequence and structural features determine constitutive and inducible zinc binding. A Primary sequence motifs of constitutive zinc binding cysteine sites. Related to Figure 4B. B Primary sequence motifs of inducible zinc binding cysteine sites. Related to Figure 4C. C Example structural folds of constitutive zinc binding sites, representative for individual structural clusters, as identified by distance analysis of the environment of constitutive zinc binding cysteines. D Example structural folds of inducible zinc binding sites, representative for individual structural clusters, as identified by distance analysis of the environment of inducible zinc binding cysteines.

**Supplementary Figure 4:**
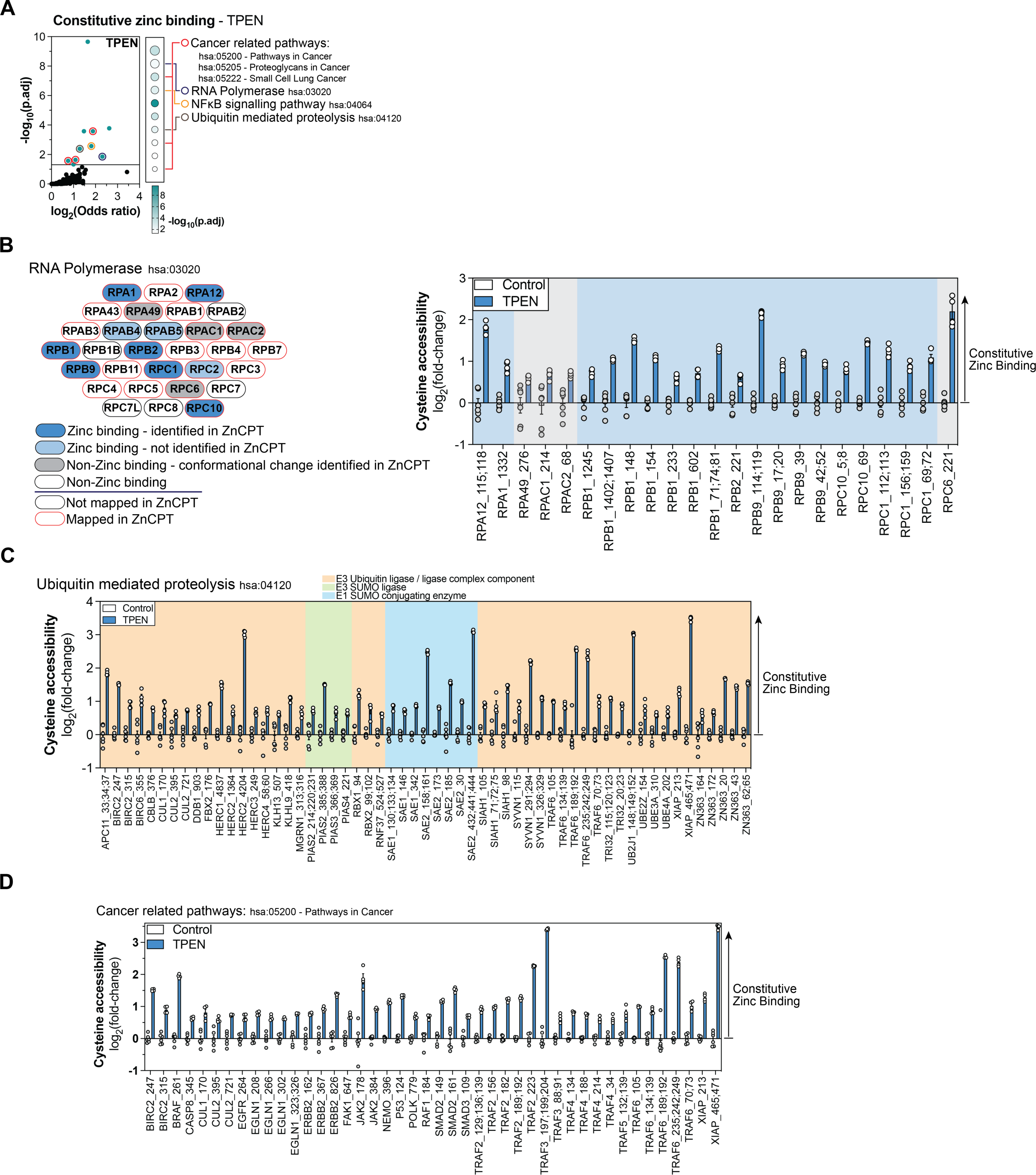
Constitutive zinc binding regulates different cellular pathways. A KEGG Pathway enrichment for constitutive zinc binding cysteines (TPEN) identifies diverse pathways as zinc-regulated. B KEGG Pathway enrichment for constitutive zinc binding cysteines (TPEN) identifies RNA Polymerase as target of zinc binding C KEGG Pathway enrichment for constitutive zinc binding cysteines (TPEN) identifies Ubiquitin E3 enzymes and SUMO E1 and E3 enzymes as targets of zinc binding D KEGG Pathway enrichment for constitutive zinc binding cysteines (TPEN) identifies several cancer-related pathways as targets of zinc binding

**Supplementary Figure 5:**
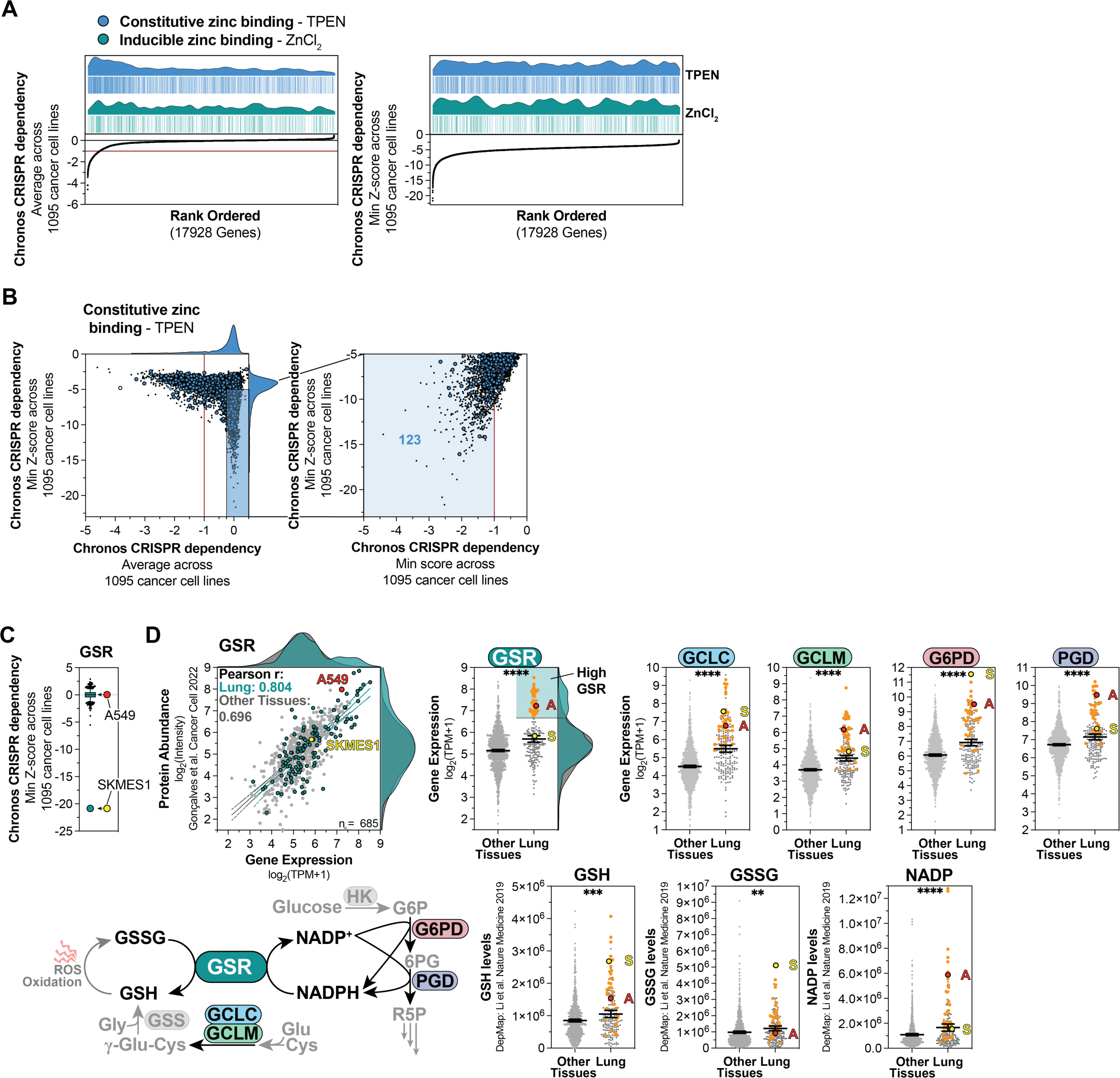
Zinc is a regulator of cancer dependency proteins with GSR being a lung specific zinc target. A Rank ordered average Chronos CRISPR dependency score and minimum Z-score for 17928 genes calculated across 1095 cancer cell lines (DepMap consortium). The distribution of constitutive (TPEN) or inducible (ZnCl_2_) zinc binding proteins identifies and enrichment of constitutive zinc binding proteins corresponding to essential genes. B Mapping the Chronos CRISPR dependency score against the minimum Z-score defines populations of essential genes and selective cell line specific dependencies amongst constitutive (TPEN) zinc binding proteins. Targets with average Chronos CRISPR dependency > −0.25 and minimum Z-score < −5 are selected and the minimum z score is mapped against the minimum Chronos CRISPR dependency score to identify strong selective dependencies (minimum Chronos CRISPR dependency < −1). C Comparison of Chronos CRISPR dependency Z-score for GSR between SKMES1 and A549 lung cancer cells (DepMap consortium) D Correlation of GSR expression and protein abundance across 685 cancer cell lines, including SKMES1 and A549 cells (DepMap consortium; Gonçalves et al. Cancer Cell 2022^59^). Select lung cancer cell lines express high levels of glutathione reductase. These cell lines also show elevated levels of glutathione synthesis genes (GCLC and GCLM), as well as genes of the pentose phosphate pathway which supply NADPH to GSR. GSR^High^ lung cancer cell lines also have elevated levels of GSH, GSSG and NADP (Li et al. Nature Medicine 2019^60^).

**Supplementary Figure 6:**
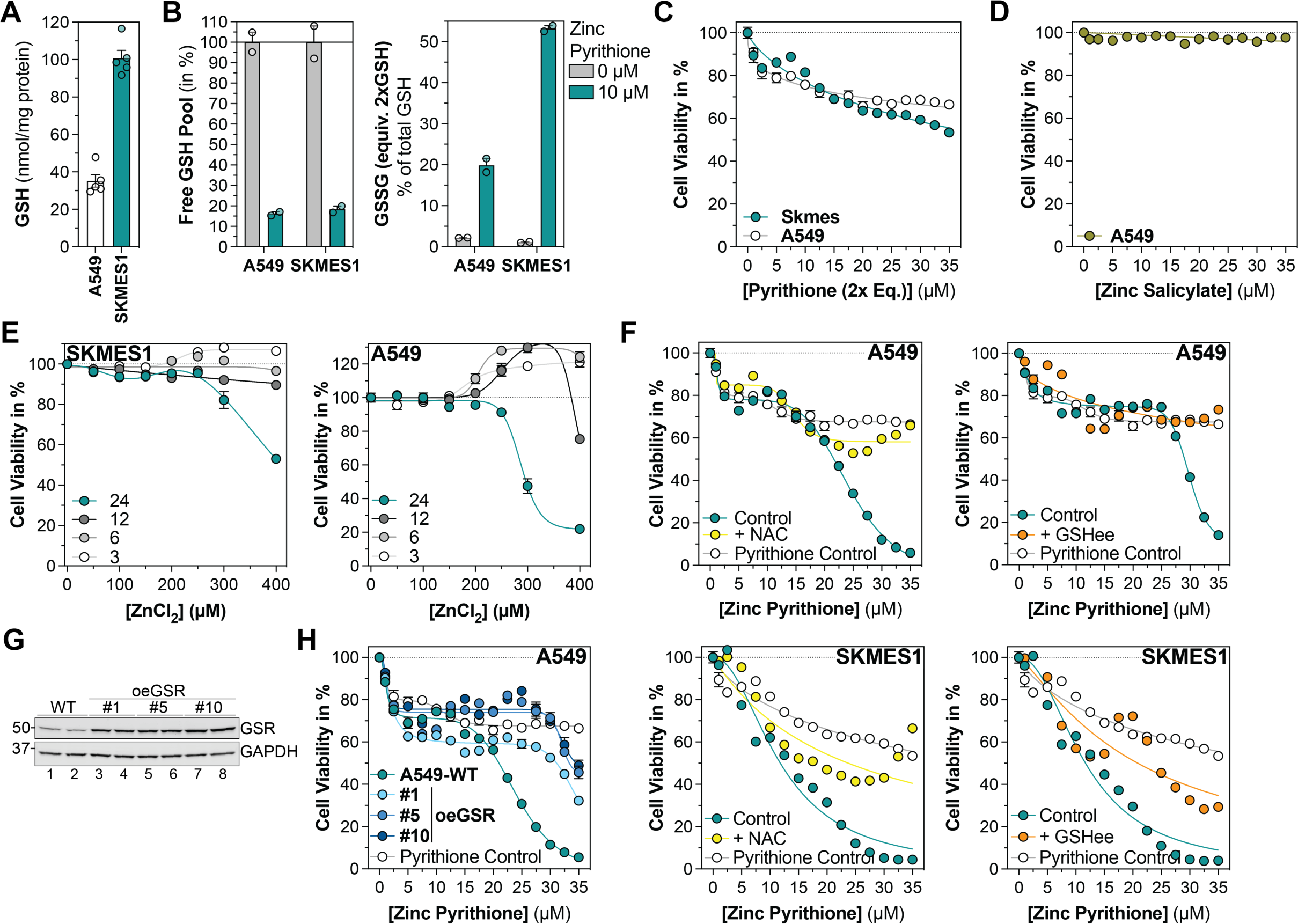
ZnCPT identifies glutathione reductase (GSR) as cancer vulnerability targetable by zinc. A Comparison of the GSH pool size between SKMES1 and A549 cancer cells B The free GSH pool of SKMES1 and A549 lung cancer cells is severely depleted, whereas the proportion of GSSG is substantially increased upon treatment with 10 µM zinc pyrithione for 24 hours. C Pyrithione inhibits cell proliferation at a concentration range at which zinc pyrithione exhibits cytotoxicity on SKMES1 and A549 cells during 24 hours of treatment. Indicated concentrations correspond to respective concentrations of zinc pyrithione, 2x equivalents of pyrithione were titrated, matching respective zinc pyrithione concentrations. D Treatment of A549 lung cancer cells with zinc salicylate does not impact cell viability during treatment for 24 hours. E Treatment of SKMES1 and A549 lung cancer cells with high concentrations of ZnCl_2_ exhibits cytotoxicity following more than 12 hours of treatment. F Treatment of SKMES1 and A549 lung cancer cells with zinc pyrithione in presence of 10 mM N-acetyl cysteine (NAC) or 1 mM Glutathione ethyl ester (GSHee) limits zinc-mediated cytotoxicity. Cell viability upon treatment with 2x equivalents of pyrithione is shown for reference. G Selected GSR overexpressing single cell clones show elevated GSR protein levels. H Overexpression of GSR in A549 lung cancer cells reduces zinc-mediated cytotoxicity upon treatment with zinc pyrithione. Cell viability upon treatment with 2x equivalents of pyrithione is shown for reference.

